# Lysosome damage triggers acute formation of ER to lysosomes membrane tethers mediated by the bridge-like lipid transport protein VPS13C

**DOI:** 10.1101/2024.06.08.598070

**Authors:** Xinbo Wang, Peng Xu, Amanda Bentley-DeSousa, William Hancock-Cerutti, Shujun Cai, Benjamin T Johnson, Francesca Tonelli, Lin Shao, Gabriel Talaia, Dario R. Alessi, Shawn M. Ferguson, Pietro De Camilli

## Abstract

Based on genetic studies, lysosome dysfunction is thought to play a pathogenetic role in Parkinson’s disease (PD). Here we show that VPS13C, a bridge-like lipid transport protein and a PD gene, is a sensor of lysosome stress/damage. Upon lysosome membrane perturbation, VPS13C rapidly relocates from the cytosol to the surface of lysosomes where it tethers their membranes to the ER. This recruitment depends on Rab7 and requires a signal at the damaged lysosome surface that releases an inhibited state of VPS13C which hinders access of its VAB domain to lysosome-bound Rab7. While another PD protein, LRRK2, is also recruited to stressed/damaged lysosomes, its recruitment occurs at much later stages and by different mechanisms. Given the role of VPS13 proteins in bulk lipid transport, these findings suggest that lipid delivery to lysosomes by VPS13C is part of an early protective response to lysosome damage.

---

Extensive genetic studies have identified many genes whose mutations cause or increase the risk of Parkinson’s disease (PD). Among these genes is VPS13C (PARK23)(*1–3*), which encodes a member of a family of bridge-like lipid transport proteins (BLTPs) that act at contacts between intracellular membranes(*4*). These are rod-like proteins with a hydrophobic groove running along their length where lipids are thought to flow unidirectionally from one bilayer to another(*5–8*). The human genome encodes four VPS13 proteins, VPS13A, VPS13B, VPS13C, and VPS13D, which have distinct, partially overlapping localizations at membrane contact sites(*9–14*), with VPS13C being localized at contacts between the endoplasmic reticulum (ER) and late endosomes/lysosomes (hence referred to as endolysosomes) as well as at ER-lipid droplet contacts (*10, 15*).

Human VPS13C is a very large protein whose approximately 30 nm long rod-like core is flanked at its C-terminal side by folded modules, namely a VAB (<u>V</u>ps13 <u>A</u>daptor <u>B</u>inding) domain, a WWE domain, an ATG2C domain (a bundle of 4 amphipathic helices so-called due to a similarity to the C-terminal region of ATG2), and a C-terminal PH domain (Fig. 1A)(*16, 17*). VPS13C binds the ER via an interaction of its N-terminal portion with the ER membrane protein VAP (an FFAT motif-dependent interaction), and endolysosomes via an interaction of its VAB domain with Rab7(*10*). It can also bind lipid droplets via its ATG2C domain. The bridge-like arrangement of VPS13C *in situ* at contacts between the ER and lysosomes has been supported by cryo-electron tomography (cryo-ET)(*17*). Building upon insights from other BLTPs, also referred to as RGB proteins because of the <u>R</u>epeating <u>β</u>-<u>G</u>roove elements that constitute their rod-like core(*7, 8*), VPS13C is hypothesized to mediate net lipid (primarily phospholipid) transport from the ER to endolysosomes(*11*). However, its precise physiological function remains elusive.

**Fig. 1.**
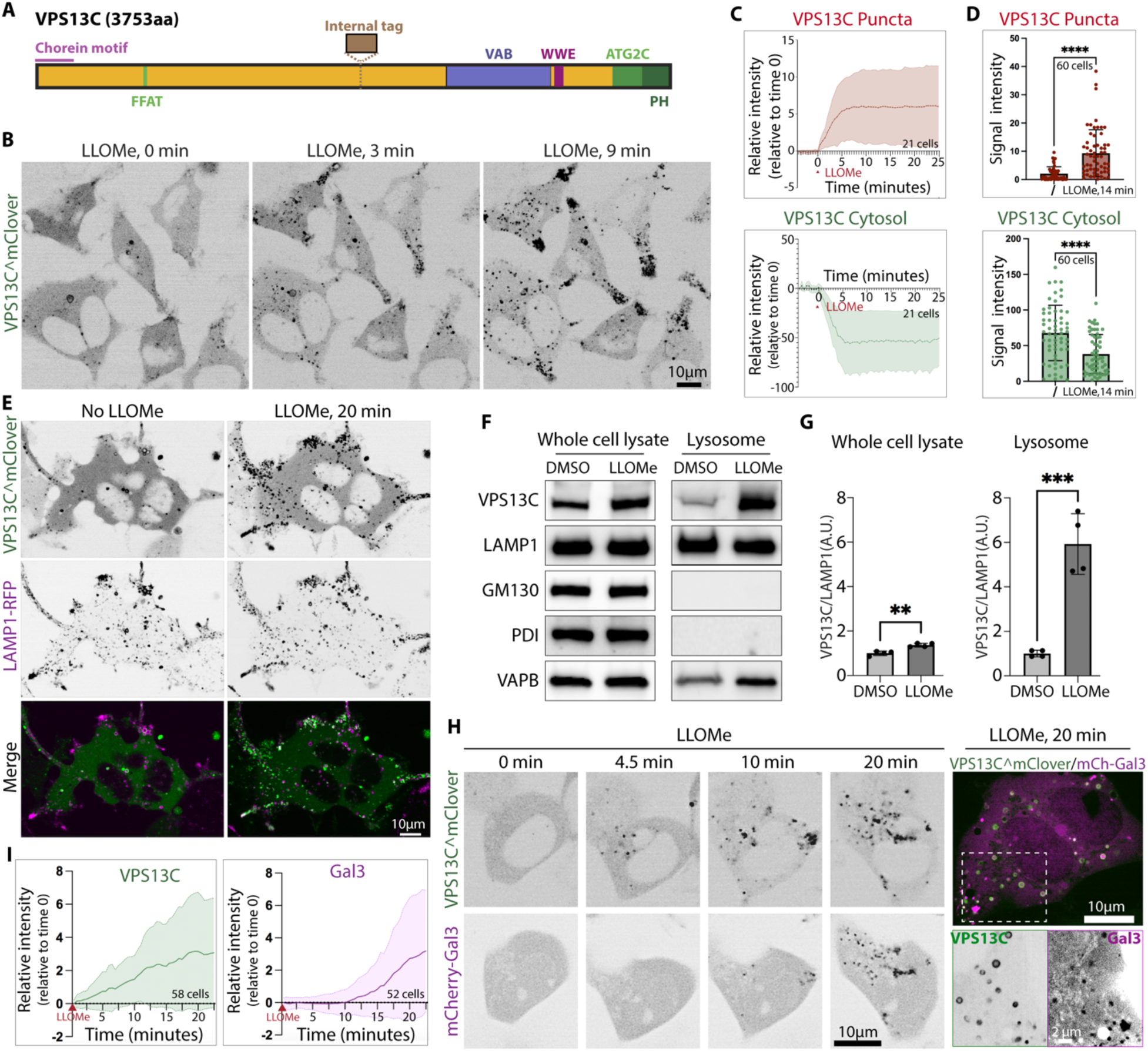
VPS13C is acutely recruited to damaged lysosomes. (A) Domain organization of human VPS13C. (B and C) Time-series of live fluorescence images of VPS13C^mClover-Flp-In cells showing rapid VPS13C recruitment to damaged lysosomes upon 1mM LLOMe treatment. Quantification of the intensity of the VPS13C^mClover punctate fluorescence or of the VPS13C^mClover cytosolic fluorescence per cell before and after 1mM LLOMe treatment from a representative experiment is shown in (C). Graphs in (C) show the normalized fluorescence relative to time 0. (D) Quantification of the intensity of the VPS13C^mClover punctate or cytosolic fluorescence per cell before and after 1 mM LLOMe treatment for 14 minutes. n = 3 biological replicates. Data were compared using two-sided t tests. Error bars represent ±SD. ****, P < 0.0001. (E) Live fluorescence images of VPS13C^mClover-Flp-In cells also co-expressing the lysosome marker Lamp1-RFP, before and after 1mM LLOMe treatment. (F and G) Western blot analysis for the proteins indicated of whole cell lysates and SPION-purified lysosomal fractions from cells treated with 1mM LLOMe or DMSO for 20 minutes. Quantification of the data is shown in (G). Bars in (G) show the normalized value relative to DMSO. Data were compared using two-sided t tests. Error bars represent ±SD. n = 4 biological replicates. **, P < 0.01; ***, P < 0.001. (H and I) Time-series of live fluorescence images of VPS13C^mClover-Flp-In cells also co-expressing mCherry-Gal3, upon 1mM LLOMe treatment. A merge field of the mClover and mCherry fluorescence at 20 min is shown at right. Quantification of the intensity of the punctate fluorescence of VPS13C^mClover or of mCherry-Gal3 before and after 1mM LLOMe treatment from a representative experiment is shown in (I). Graphs in (I) show the normalized fluorescence relative to time 0. n = 3 biological replicates. All the individual channel images in this figure are shown as inverted grays.

Loss-of-function mutations in VPS13C cause PD (*1*). Previous studies demonstrated that the genetic deletion of VPS13C in HeLa cells or iPSC-derived human neurons results in altered lysosome homeostasis(*15, 18*). In these cells, TFEB, a master regulator of lysosome biogenesis, is activated and the number of lysosomes is increased, most likely reflecting compensatory mechanisms to cope with lysosomal dysfunction(*15*). Notably, in VPS13C KO HeLa cells the cGAS-STING innate immunity pathway(*19*) is activated potentially reflecting impaired STING degradation in dysfunctional lysosomes and escape of mitochondrial DNA into the cytosol(*15*). These findings indicate a potential role for VPS13C in maintaining the integrity of the lysosomal membrane, and possibly also in membrane repair. They are especially relevant in the context of PD, as several PD-related genes encode proteins implicated in lysosomal function(*20–22*). These include the gene encoding leucine-rich repeat kinase 2 (LRRK2), the most frequently mutated protein in familial PD(*23, 24*). Despite its predominant cytosolic distribution, LRRK2 can be acutely recruited to lysosomes in response to various manipulations that induce perturbation of their membranes, including addition to cells of L-leucyl-L-leucine methyl ester (LLOMe), a lysosomotropic agent that accumulates in lysosomes and is converted to membranolytic metabolites in the lysosome lumen(*25, 26*). Several factors other than LRRK2 are recruited to endolysosomes in response to damage. These include ESCRT proteins that function as key players in membrane repair mechanisms(*27–31*). They also include the ER-anchored OSBP/ORP family lipid transport proteins that are recruited in response to the generation of phosphatidylinositol-4-phosphate (PI4P) on endolysosome membranes in the so-called PITT pathway(*32–34*).

While investigating VPS13C localization and dynamics in a variety of cultured cells, we observed that this protein was only present at a subset of endolysosomes, suggesting the requirement of specific conditions for such localization. Prompted by these findings, we investigated whether the recruitment of VPS13C and the formation/expansion of VPS13C-dependent contacts between the ER and lysosomes are part of a cellular response to lysosome damage. We found that formation of VPS13C-dependent ER-endolysosome tethers is part of an early response to lysosome damage and is differentially controlled relative to the recruitment of ESCRT, OSBP/ORP family proteins and LRRK2. Absence of VPS13C makes lysosomes more sensitive to LLOMe-induced perturbation. Our results also show that localization of VPS13C at the interface between the ER and lysosomes requires binding of its ATG2C domain to the damaged lysosome membrane leading to the release of an inhibition that prevents access of VPS13C to Rab7 on the surface of endolysosomes.

## RESULTS

### VPS13C is acutely recruited to damaged lysosomes

To track the localization of VPS13C within cells, we used Flp-In TREx 293 cells to generate a stable cell line expressing VPS13C tagged with mClover at an internal site (VPS13C^mClover) under the control of tetracycline (Fig. 1A and fig. S1A). Anti-VPS13C western blotting of homogenates of these cells revealed that, under the conditions used for our experiments (24 or 48 hours in the presence of 0.1 µg/ml tetracycline), no global overexpression of VPS13C occurred as the levels of endogenous VPS13C were reduced in parallel with expression of exogenous VPS13C^mClover (Fig. S1B).

Confocal imaging of these VPS13C^mClover expressing cells, defined henceforth as VPS13C^mClover-Flp-In cells, showed puncta or doughnut-like structures (Fig. 1B), which corresponded to a small subset of lysosomes, as confirmed by colocalization with the lysosome marker LAMP1-RFP (Fig. 1E). An additional pool of VPS13C was diffusely distributed throughout the cytosol. However, upon addition of 1 mM LLOMe, a lysosomotropic agent that causes lysosomal membrane perturbation(*35*), a rapid (within minutes) and massive recruitment of VPS13C^mClover to the majority of LAMP1-RFP-positive organelles was observed (Fig. 1B-E). Notably, LAMP1-positive lysosomes were concentrated at the tips of cells processes in this cell line and their localization was not affected by exposure to LLOMe (Fig. 1E). Recruitment of VPS13C to lysosomes correlated with lysosomal membrane damage, as pre-incubating cells with the cathepsin C inhibitor E64d, the enzyme that processes LLOMe into membranolytic polymers, inhibited VPS13C recruitment induced by LLOMe treatment (Fig. S1C).

To validate with a biochemical approach the recruitment of VPS13C to lysosomes in response to LLOMe and to determine whether such recruitment is also observed for endogenous untagged VPS13C, we used a previously described technique for the purification of lysosomes. The method is based on incubation of live cells with dextran-conjugated superparamagnetic iron oxide nanoparticles (SPIONs) to allow endocytic uptake followed by a chase to allow traffic of the particles to lysosomes, and subsequent cell lysis and recovery of lysosomes on a magnetic column(*15, 36, 37*). Western blot analysis of the magnetically isolated material revealed abundant presence of LAMP1, but not of GM130 (Golgi marker) and PDI (a luminal ER marker), confirming the successful and selective isolation of lysosomes (Fig. 1F). Importantly, immunoblotting for VPS13C showed a very robust enrichment of the endogenously expressed VPS13C protein relative to LAMP1 in cells treated with LLOMe (Fig. 1F, quantified in Fig. 1G) confirming the imaging experiments. A slight increase in the ER protein VAP is in line with the recruitment of VPS13C (Fig. 1F), as VPS13C binds VAP and fragments of ER membranes are expected to copurify with endolysosomes.

### The recruitment of VPS13C to damaged lysosomes precedes the recruitment of galectin 3 and also of ESCRT-III

Expression of fluorescently-tagged galectin3 has been used as a tool to detect severely damaged lysosomes(*27, 38*). Galectin 3 is a β-galactoside binding lectin that remains soluble in the cytosol of healthy cells. However, it rapidly translocates to damaged lysosomes, where it binds the glycoprotein-rich luminal surface of their membranes(*38*). When transiently expressed in VPS13C^mClover-Flp-In cells, mCherry-tagged galectin3 (mCherry-Gal3) had a cytosolic localization (as expected) under basal conditions. LLOMe treatment induced Gal3 puncta formation, indicating its recruitment to damaged lysosomes (Fig. 1H). Importantly, however, recruitment of VPS13C^mClover preceded the recruitment of galectin 3 to lysosomes by several minutes (Fig. 1H and I) implying that VPS13C senses a change in the properties of the lysosomal membrane that precedes severe membrane rupture.

An early response to lysosome damage by LLOMe, which also precedes galectin 3 recruitment, is the recruitment of the ESCRT complex (*27, 28*). Comparison of the LLOMe-induced recruitment of VPS13C (VPS13C^mClover) and of the ESCRT-III component IST1 (IST1-Apple) suggested an earlier recruitment of VPS13C relative to IST1 (Fig. S2A). Moreover, while the recruitment of IST1 triggered by LLOMe was inhibited by the chelation of cytosolic Ca^2+^ by BAPTA, consistent with such recruitment being dependent on the leakage of Ca^2+^ from damaged lysosomes (*27, 39, 40*), the recruitment of VPS13C was not blocked by BAPTA (Fig. S2B). Thus, although both the recruitment of VPS13C and of ESCRTIII represent a very early response to lysosome damage, their recruitment appears to be mediated by different mechanisms.

### VPS13C recruited to lysosomes is bound to the ER

The proposed bridge-like lipid transport function of VPS13C at ER-endolysosome contacts implies its simultaneous binding to both the ER and endolysosomes. Binding to the ER relies on the interaction of its FFAT motif with the transmembrane ER protein VAP (VAPA and VAPB), while binding to the endolysosomal membrane requires the interaction of its VAB domain with Rab7(*10, 15*) (Fig. 2A). If LLOMe-induced recruitment of VPS13C to lysosomes is aimed at allowing a flux of lipids from the ER to damaged membrane of these organelles, the recruitment of VPS13C to lysosomes should be accompanied by the recruitment to these organelles of the ER via VAP. Consistent with this hypothesis, following LLOMe treatment of VPS13C^mClover-Flp-In cells that additionally express mCherry tagged VAPB (mCherry-VAPB), the recruitment of VPS13C was accompanied by the enrichment of VAP at endolysosomes (Fig. 2B).

**Fig. 2.**
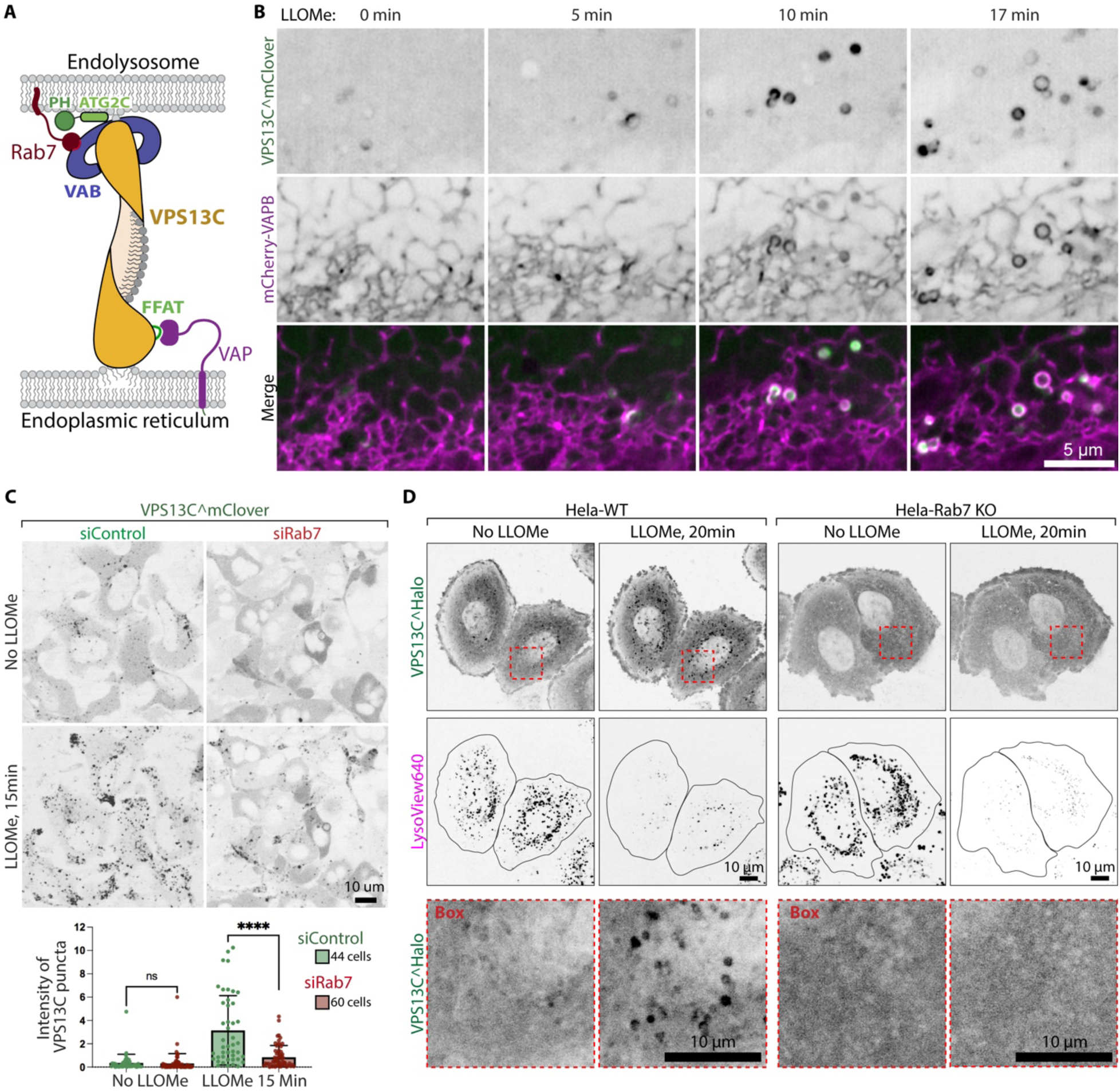
VPS13C functions at the ER-endolysosome membrane contact sites. (A) Cartoon depicting VPS13C localized at the ER-endolysosome membrane contact sites. (B) High-magnification of time-series images of VPS13C^mClover-Flp-In cells also co-expressing exogenous mCherry-VAPB, upon 1mM LLOMe treatment. (C) Live fluorescence images of VPS13C^mClover-Flp-In cells showing VPS13C^mClover localization before and after 1mM LLOMe treatment in control and Rab7 knockdown cells. Quantification of fluorescence intensity of VPS13C^mClover puncta signals per cell before and after LLOMe treatment is shown at the bottom. Data were compared using two-sided t tests. n = 3 biological replicates. Error bars represent ±SD. (D) Live fluorescence images of WT or Rab7 KO Hela cells expressing VPS13C^Halo and labeled with LysoView 640 (a luminal marker of acidic lysosomes) before and after 1mM LLOMe treatment. Boxed regions from the VPS13C^Halo channel is shown at the bottom. All the individual channel images in this figure are shown as inverted grays.

### Rab7 dependence of the VPS13C recruitment

Binding of VPS13C to lysosomes was shown to be dependent on the interaction of its VAB domain with Rab7 and to be abolished by expression of dominant negative Rab7(*15*). To explore whether such interaction is required for the acute recruitment of VPS13C to lysosomes in response to LLOMe, Rab7 was knocked down by siRNA in VPS13C^mClover-Flp-In cells (fig. S3A). The accumulation of VPS13C at lysosomes in response to LLOMe was significantly reduced in Rab7 knockdown cells compared to WT cells (Fig. 2C). To further validate this observation, we expressed VPS13C^Halo in previously generated Rab7 KO HeLa cells(*41*), which were also incubated with LysoView 640 to visualize lysosome. In these cells, puncta of VPS13C fluorescence reflecting accumulation of VPS13C at lysosomes no longer occurred either under basal conditions or after LLOMe treatment (Fig. 2D). Occurrence of lysosome damage after LLOMe was confirmed by the loss of LysoView640 signal. Interestingly, the protein level of VPS13C was significantly elevated in Rab7 KO cells (Fig. S3B), possibly as a compensatory response to lysosomal stress induced by the absence of Rab7. We thus conclude that not only basal binding of VPS13C to lysosomes, but also its enhanced recruitment upon lysosome damage is a Rab7-dependent event. These findings prompted us to examine how Rab7-dependent recruitment of VPS13C is regulated.

### VPS13C recruitment to endolysosomes is not dependent on Rab7 phosphorylation state

Phosphorylation of Rab GTPases has emerged as an important factor in the regulation of their functions(*42–44*). Specifically, phosphorylation of Rab7 at Ser72 functions as a phosphoswitch that determines its interactions with distinct downstream effectors(*45, 46*). Tank-binding kinase 1 (TBK1) and leucine-rich repeat kinase 1 (LRRK1) mediate this phosphorylation event, while its dephosphorylation is regulated by the phosphatase PPM1H(*45, 47, 48*). As shown by Fig 3A, we found that Rab7 undergoes fast and robust Ser72 phosphorylation in VPS13C^mClover-Flp-In cells upon LLOMe treatment, with the time course of this event closely mirroring that of VPS13C recruitment shown in Fig.1. To investigate potential contributions of LRRK1 and TBK1 to this LLOMe-dependent increase of Rab7 phosphorylation at Ser72, we assessed the impact of these kinases in previously generated KO mouse embryonic fibroblasts (MEFs)(*47*) and found that such increase was blocked by the KO of LRRK1 but not by the double KO of TBK1 and of its paralogue IKKe (fig. S4). On the other hand, in HeLa cells, TBK1, which can also be activated by LLOMe (*49*), was found to be responsible for ∼50% of the increase in Rab7 Ser72 phosphorylation induced by LLOMe(*41*), suggesting the occurrence of cell-specific differences in this event.

**Fig. 3.**
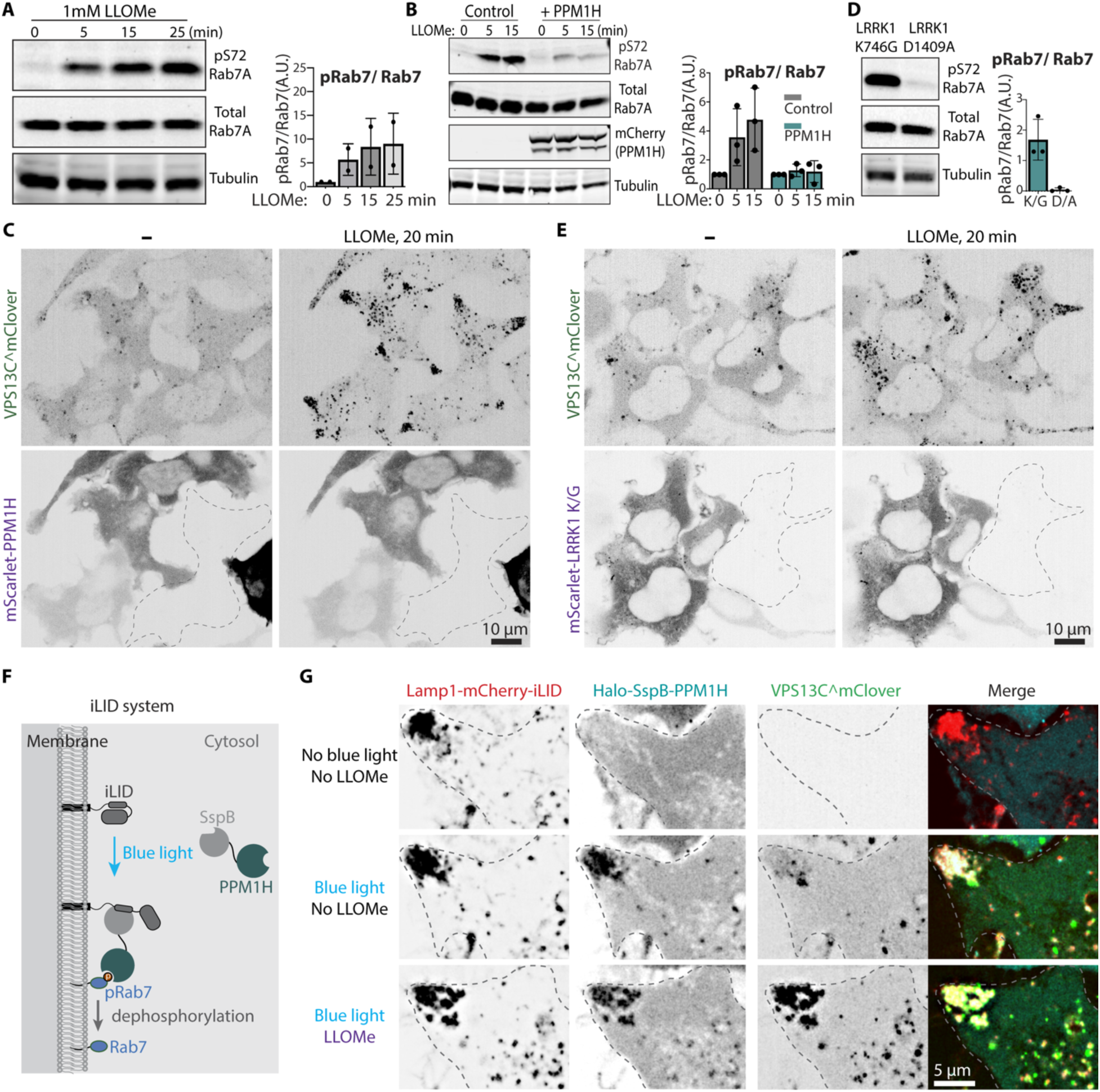
Rapid Rab7 phosphorylation induced by LLOMe is unnecessary for VPS13C recruitment. (A and B) Western blot analysis for the proteins indicated of whole cell lysates from VPS13C^mClover-Flp-In cells at various time after treatment with 1mM LLOMe. A sample of cells in (B) were transfected with mScarlet-PPM1H. Quantification of the data is shown on the right. Bars show normalized values relative to time 0. n = 2 biological replicates for (A). n = 3 biological replicates for (B). (C) Live fluorescence images of VPS13C^mClover-Flp-In cells also expressing mScarlet-PPM1H before and after 1mM LLOMe treatment. Cells not expressing mScarlet-PPM1H are indicated by dashed lines. (D) Western blots for total Rab7 and S72 phospho-Rab7 (and of tubulin as a loading control) of whole cell lysates from VPS13C^mClover-Flp-In cells also co-expressing constitutively kinase active (mScarlet-LRRK1^K746G^) or kinase dead (mScarlet-LRRK1^D1409A^) LRRK1 mutant. Quantification of the data is shown on the right. Bars show normalized values relative to the signal obtained with kinase dead LRRK1 (D1409A). n = 3 biological replicates. (E) Live fluorescence images of VPS13C^mClover-Flp-In cells also expressing constitutively kinase active LRRK1 mutant (mScarlet-LRRK1^K746G^) before and after 1mM LLOMe treatment. Cells not expressing mScarlet-LRRK1 ^K746G^ are indicated by dashed lines. (F) Schematic drawing depicting the iLID light-dependent protein heterodimeric system used in (G). (G) Live fluorescence images of VPS13C^mClover-Flp-In cells also co-expressing exogenous Lamp1-mCherry-iLID (bait) and Halo-SspB-PPM1H (prey), before and after blue light illumination, as well as before and after 1mM LLOMe treatment. All the individual channel images in this figure are shown as inverted grays.

To directly test the role of Rab7 Ser72 phosphorylation to the regulation of VPS13C recruitment, we next implemented multiple strategies to target this post-translational modification. First, we transiently overexpressed PPM1H(*47*) into VPS13C^mClover-Flp-In cells to retain Rab7 in its dephosphorylated state. Western blot analysis confirmed a significant reduction of Rab7 phosphorylation after LLOMe treatment (Fig. 3B), but the recruitment of VPS13C remained unaffected in cells expressing PPM1H compared to their PPM1H-negative neighbor cells (Fig. 3C). Conversely, overexpression of a constitutively kinase-active LRRK1 mutant, LRRK1^K746G^ (but not the kinase dead LRRK1^D1409A^ mutant)(*47*), robustly induced Rab7 Ser72 phosphorylation under basal conditions (Fig. 3D), but failed to increase the recruitment of VPS13C under either basal conditions or following LLOMe treatment (Fig. 3E). To exclude the possibility that these negative results may reflect adaptive changes upon expression of PPM1H or mutant LRRK1, we explored the effect of acute manipulation of Rab7 phosphorylation at the lysosomal surface. To this aim we employed the improved(*50*) light-induced dimerization (SspB, iLID) system(*51*) to recruit PPM1H to lysosome membranes via blue light (Fig. 3F). We observed rapid translocation of cytosolic SspB-linked PPM1H to lysosomes upon blue light illumination (Fig. 3G, middle), but this translocation did not prevent the prompt recruitment of VPS13C following LLOMe treatment (Fig. 3G, bottom). Altogether, while these results demonstrate a very robust increase in Rab7 phosphorylation upon LLOMe treatment, such phosphorylation reaction is not required for VPS13C recruitment.

### The access of full length VPS13C to Rab7 on endolysosomes undergoes regulation

As mentioned above, our previous findings indicate that binding of VPS13C to lysosomes depends on the interaction of its VAB domain with Rab7(*10, 15*). While VPS13C recruitment to lysosomes by LLOMe could be explained by an enhanced activated state of Rab7, a discrepancy that we have observed between the localization of the VAB domain fragment of VPS13C and the localization of full-length VPS13C speaks against this possibility. As shown in Fig.1E, before LLOMe addition, VPS13C was enriched only on a small subset of lysosomes in VPS13C^mClover-Flp-In cells. In contrast, when the VAB domain (Fig. 4A) tagged with mCherry (mCh-VAB), which mediates the interaction with Rab7, was expressed in these cells, a surprising difference was observed. Robust colocalization of mCh-VAB with the great majority of organelles positive for the lysosomal marker LysoView 640 (a probe that accumulates within acidic lysosomes) occurred even under basal conditions (Fig. 4B left). This massive localization of the VAB domain on lysosomes was further validated in HeLa cells (Fig. S5A). Importantly, this association of the VAB domain with lysosomes was lost in Rab7 knockout HeLa cells, further confirming the constitutive binding of this domain to Rab7 in the absence of any external manipulation (Fig. S5A). Minutes after LLOMe treatment, when loss of LysoView 640 signal from most lysosomes occurred as a result of their damage, much of the VAB domain was shed and this coincided with the accumulation on lysosomes of the full-length VPS13C^mClover with a reduction of its cytosolic pool (Fig. 4B left). Interestingly, the few lysosomes that remained strongly positive for the LysoView640 signal even after LLOMe treatment also retained a strong VAB signal and failed to recruit full-length VPS13C^mClover (Fig. 4C). These results indicate that access to Rab7 of full-length VPS13C, but not of its VAB domain alone, is regulated.

**Fig. 4.**
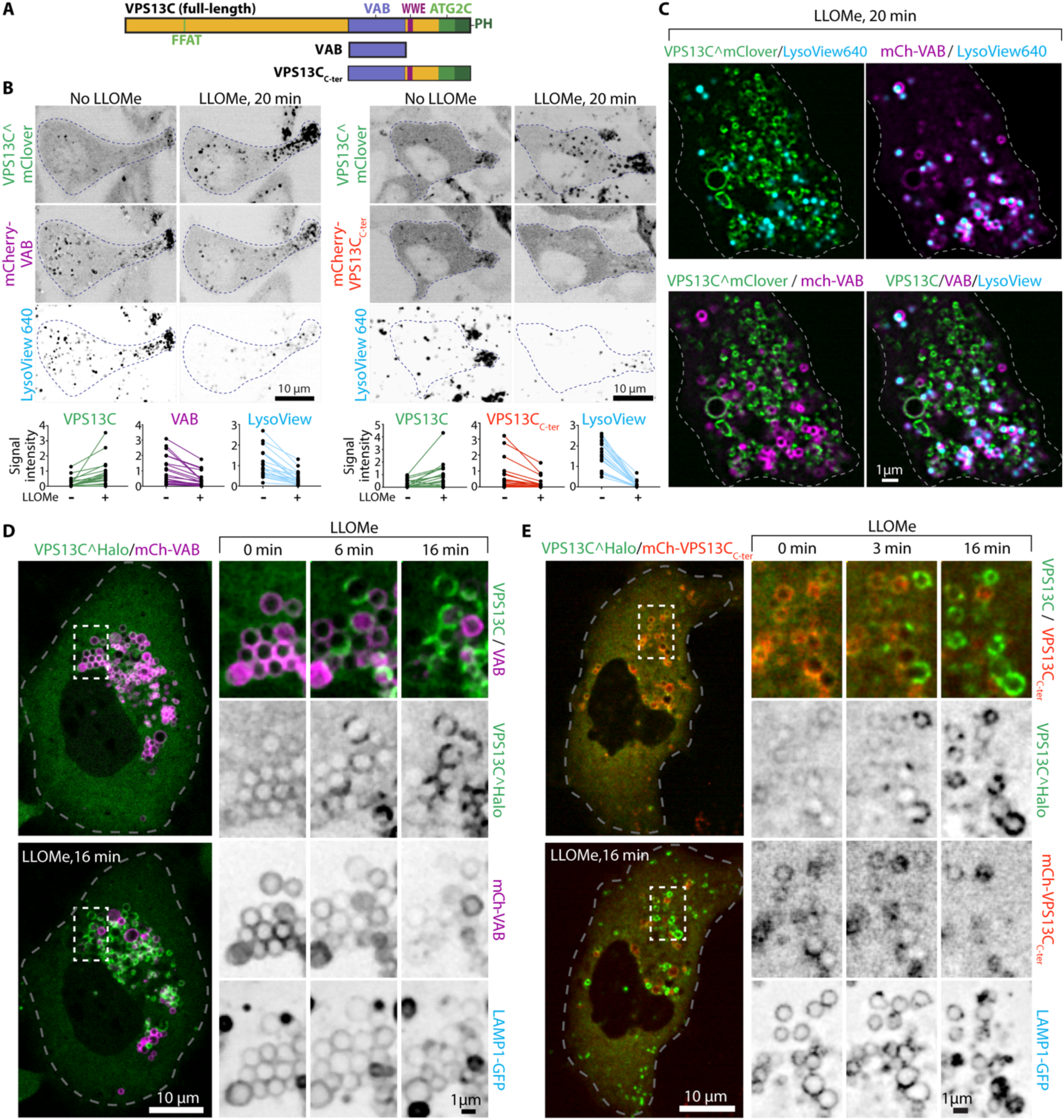
An intramolecular regulation controls access of full length VPS13C to Rab7. (A) Domain organization of full-length VPS13C. Deletion constructs used for the experiments of this figure are also indicated. (B) Live fluorescence images of VPS13C^mClover-Flp-In cells also co-expressing the mCherry-VAB domain (left) or mCherry-VPS13C_C-ter_ (right) and labeled with LysoView 640 (a luminal marker of acidic lysosomes) before and after 1mM LLOMe treatment. Quantification of the punctate fluorescence from individual channels in a representative experiment is shown at the bottom. Each line represents the average intensity of the indicated signals from the same cell before and after LLOMe treatment. 20 cells were analyzed. (C) High-magnification of live fluorescence images of VPS13C^mClover-Flp-In cells also expressing mCherry-VAB, 20 minutes after addition of 1mM LLOMe. Note almost no overlap between mCherry-VAB and VPS13C^mClover fluorescence. (D and E) Live fluorescence images of RPE1 cells co-expressing exogenous VPS13C^Halo and either mCherry-VAB (D) or mCherry-VPS13C_C-ter_ (E) before and after 1mM LLOMe treatment. Time-series of individual channels from the boxed regions are shown on the right in both (D) and (E).

To further explore this regulation, we monitored the dynamics of co-expressed full-length VPS13C and VAB domain in hTERT RPE-1 (RPE1) cells, i.e. cells where the presence of large lysosomes facilitates the tracking of individual vesicles over time. Even in these cells, strong lysosomal binding of the VAB domain, but only scattered lysosomal binding of full-length VPS13C, was observed under basal conditions (Fig. 4D), while a swap occurred after LLOMe treatment, with an increased lysosome association of the full-length protein and a decrease association of the VAB domain (Fig. 4D right). As binding to Rab of Rab effectors reflects a dynamic equilibrium of binding/unbinding reactions, one potential explanation for this finding is that the recruitment of the ER (Fig. 2B) to the lysosomal membranes creates a barrier that prevent rebinding of the VAB domain of VPS13C once the ER has covered the endolysosomal membrane. Not only endogenous VPS13C, but also other ER-lysosomal tethers were recently shown to be recruited to lysosomes in response to LLOMe(*32, 33*), and could thus contribute to this ER recruitment-dependent effect. In fact, a decrease in VAB binding upon LLOMe treatment did not require overexpression of VPS13C (fig. S5B left). Collectively, these findings suggest that the availability of active (GTP loaded) Rab7 is not limiting, and that in full-length VPS13C the accessibility of the VAB domain to Rab7 binding may be subject to a LLOMe-regulated mechanism.

### Evidence for an intramolecular regulation of VPS13C binding to Rab7 on lysosomes

One factor that could limit accessibility to lysosome-associated Rab7 of full-length VPS13C in the absence of lysosome damage is an intramolecular interaction blocking the Rab7 binding site in VPS13C (Fig. 4A). The VAB domain of VPS13C is flanked by other folded modules (Fig. 2A), a WWE domain, the ATG2C domain and a PH domain, which project out of the rod-like core of VPS13C(*17*). The position of these modules relative to each other and to the rod-like core is not fixed, as suggested by low resolution negative staining EM analysis of yeast VPS13(*52*). To test a potential inhibitory interaction between the VAB domain and other folded modules that flank the rod-like core of VPS13C at its C-terminal region, we expressed in VPS13C^mClover-Flp-In cells the mCherry-tagged C-terminal fragment of VPS13C (mCh-VPS13C_C-ter_), which comprises the VAB, WWE, ATG2C, and PH domains, in addition to a small fragment of the rod-like core (Fig. 4A and B right). We found that the VPS13C_C-ter_ fragment acted similarly to full-length VPS13C, with few lysosomes positive for this construct and a predominant cytosolic localization under basal conditions (Fig. 4B right and Fig. S5C left). These results support the hypothesis that some structural element within the C-terminal fragment of VPS13C obstructs access of the VAB domain to Rab7. However, in contrast to what was observed with full length VPS13C, even this low binding to lysosomes observed under basal conditions was mostly lost upon exposure to LLOMe, when robust recruitment of full-length VPS13C was observed (Fig. 4B right). Similar results were observed when VPS13C^Halo and the mCherry-VPS13C_C-ter_ construct were co-expressed in RPE1 cells (Fig. 4E and fig. S5B, C right). These results support the hypothesis that recruitment of the ER to lysosomes reduces the association with their surface of cytosolic factors bound to Rab7 by hindering their access to such surface.

Based on these findings we suggest that an intramolecular interaction within the C-terminal fragment of VPS13C prevents the binding of the VAB domain to Rab7 and that a signal(s) generated by lysosomal damage is required to release such inhibition.

### The ATG2C domain of VPS13C detects lysosome membrane damage

To directly test whether the C-terminal fragment of VPS13C, which contains the ATG2C and PH domains (Fig. 1A), blocks VAB domain access for Rab7 binding, we generated a VPS13C deletion construct lacking these domains, VPS13C-Δ(ATG2C-PH) (Fig. 5A). Expression of this construct with an internal Halo tag, VPS13C-Δ(ATG2C-PH)^Halo, in RPE1 cells revealed strong lysosomal binding (Fig. 5B) under basal conditions, comparable to the binding of VAB domain alone (Fig. 4D). This was in contrast to the faint lysosomal binding observed for the full-length VPS13C^Halo under these conditions (Fig. 5B and Fig. 4D, E). We conclude that the C-terminal fragment of VPS13C, which comprises the ATG2C and PH domains, acts as a brake that prevents the VAB domain from accessing Rab7 in the absence of lysosome damage (Fig. 5C).

**Fig. 5.**
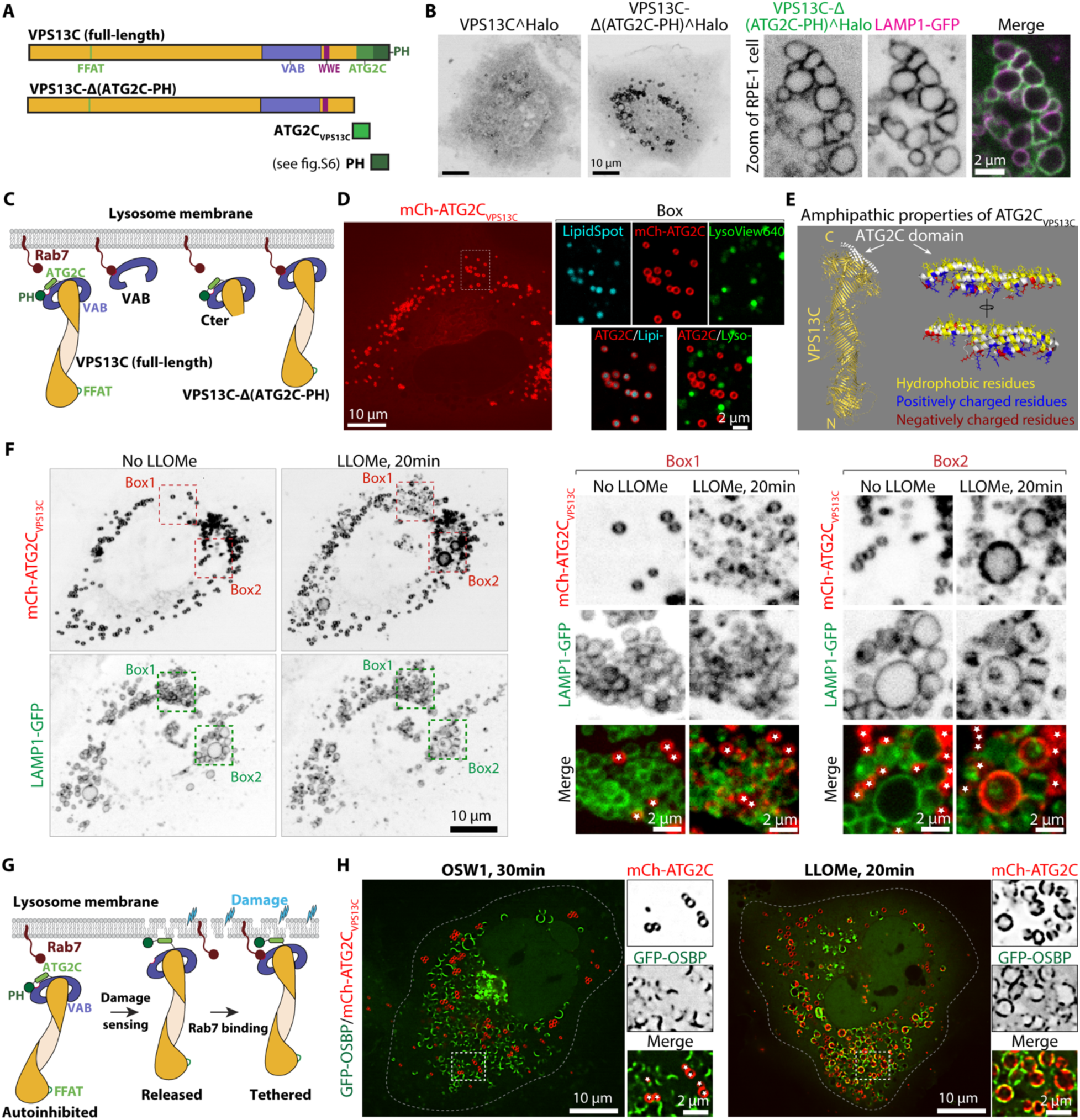
The ATG2C domain of VPS13C detects lysosome membrane damage. (A) Domain organization of either full-length VPS13C or domains of VPS13C used for the experiments of this figure. (B) Live fluorescence images of RPE1 cells expressing full-length VPS13C^Halo or VPS13C-Δ(ATG2C-PH)^Halo under basal conditions. Zoom of a region from RPE1 cells on the right showing co-localization of VPS13C-Δ(ATG2C-PH)^Halo with Lamp1-GFP. (C) Cartoon depicting the proposed association of VPS13C and its deletion constructs with Rab7 on the surface of lysosomes. (D) Live fluorescence images of RPE1 cells expressing mCherry-ATG2C_VPS13C_ and labeled with Lysoview 640 and LipidSpot 488 (a marker of lipid droplet). (E) Predicted structure of the ATG2C domain of VPS13C based on AlphaFold3 showing at right high-power views of the amphipathic helices. (F) Live fluorescence images of RPE1 cells expressing mCherry-ATG2C_VPS13C_ before and after 1mM LLOMe treatment. Boxed regions are shown at higher magnification on the right demonstrating co-localization of mCherry-ATG2C_VPS13C_ with Lamp1-GFP after LLOMe treatment. White stars label lipid droplets. (G) Putative model illustrating how binding of the ATG2C_VPS13C_ domain to the bilayer may release an autoinhibitory conformation of VPS13C to allow its binding to Rab7 on the lysosomal surfaces. (H) Live fluorescence images of RPE1 cells expressing mCherry-ATG2C_VPS13C_ and GFP-OSBP after 1mM LLOMe (left) or 20nM OSW1 (right) treatment. White stars label lipid droplets.

To further explore how the ATG2C-PH domain portion of VPS13C controls its binding to lysosomes, we expressed mCherry-tagged versions of the ATG2C domain (mCh-ATG2C_VPS13C_) and of the PH domain (mCh-PH) separately in RPE1 cells (Fig. 5A, D, and fig. S6A). Under basal conditions, mCh-PH predominantly localized to the cytosol (Fig. S6A), while mCh-ATG2C_VPS13C_ exhibited strong lipid droplet binding (Fig. 5D), as previously reported(*10*). This binding is consistent with the amphipathic nature of the 4 parallel helices that constitute this domain, as lipid droplet binding is a feature of many amphipathic helices(*53, 54*) (Fig. 5E). Following LLOMe treatment, an additional appearance of an mCh-ATG2C_VPS13C_-positive signal that overlapped with LAMP1 was observed (Fig. 5F), while mCh-PH remained primarily cytosolic (Fig. S6A). These findings suggest that the ATG2C_VPS13C_ domain can detect damaged lysosome membranes and bind to them.

Altogether, these results support a model according to which under basal conditions intramolecular interactions involving the C-terminal ATG2C_VPS13C_ domain and/or PH domain prevent the VAB domain from accessing Rab7. Upon lysosomal membrane damage, binding of the ATG2C_VPS13C_ domain to these membranes results in a conformational change of the C-terminal moiety of VPS13C which releases the VAB domain and enables its interaction with Rab7, thereby further promoting and stabilizing the interaction of VPS13C with lysosomes (Fig. 5G).

### The PITT pathway is not involved in VPS13C recruitment

The acute recruitment of VPS13C to lysosomes is not the only event leading to the formation of ER-lysosome tethers in response to lysosome damage. It was reported that exposure to LLOMe results in a rapid accumulation of phosphatidylinositol-4-phosphate (PI4P) on their surface via the recruitment of PI4K2A. PI4P at the lysosome surface, in turn, recruits ER-anchored oxysterol-binding protein (OSBP)-related proteins (ORP) family members, i.e. shuttle-type lipid transfer proteins which mediate transport of PI4P to ER in exchange for countertransport of PtdSer and cholesterol from the ER to lysosomal surface, possibly helping repair lysosome damage(*32–34*). The timing of this response to lysosome damage, referred to as phosphoinositide-initiated membrane tethering and lipid transport (PITT) pathway(*32*), is similar to the timing of VPS13C recruitment to lysosomes (occurring within minutes after LLOMe treatment), prompting us to investigate whether PI4P contributes to the recruitment of the ATG2C domain of VPS13C. To do so, we evaluated in RPE1 cells the impact of OSW-1(*55*) on the recruitment of mCh-ATG2C_VPS13C_ to lysosomes. OSW1 is a natural compound that blocks the PI4P/cholesterol counter-transfer function of OSBP and thus robustly enhances presence of PI4P on lysosomes. In agreement with previous observations(*56–58*), we found that under basal conditions GFP-OSBP was exclusively localized in the Golgi complex region (fig. S6B), while after treatment of cells with OSW-1 a pool of GFP-OSBP rapidly translocated to lysosomes (Fig. S6B), consistent with PI4P accumulation on their membranes, while no obvious translocation of ATG2C_VPS13C_ to these organelles occurred (Fig. 5H, and fig. S6B). Both GFP-OSBP and ATG2C_VPS13C_, however, could be recruited to lysosome in response to LLOMe-induced damage (Fig. 5H, and Fig. S6C)). Thus, ATG2C_VPS13C_ is not a PI4P effector. Similar results were obtained with full-length VPS13C, as in VPS13C^mClover-Flp-In cells, OSBP translocated to lysosomes in response to OSW-1 but full-length VPS13C did not (fig. S7A). This further supports the conclusion that the recruitment of VPS13C to damaged lysosomes is independent of PI4P. Additionally, while both VPS13C and OSBP were rapidly recruited to lysosomes upon LLOMe treatment, the recruitment of VPS13C occurred earlier than the recruitment of OSBP (Fig. S7B). We conclude that while both VPS13C and OSBP detect LLOMe-induced lysosome damage, the mechanisms of their recruitment are different and that the main trigger for VPS13C binding to damaged lysosomes is a perturbation of lipid packing in their membrane (see Discussion).

Finally, although the recruitment of VPS13C to lysosomes in response to LLOMe slightly preceded the recruitment of OSBP and IST1, we found that both proteins were still efficiently recruited to lysosomes following LLOMe treatment in VPS13C KO A549 cells (Fig. S8A-C). This finding indicates that lysosomal recruitment of OSBP and IST1 is independent of VPS13C.

### Loss of VPS13C increases the fragility of lysosomes

The rapid response of VPS13C to lysosomal damage, along with its lipid transfer properties, suggest that VPS13C may be important to preserve the integrity of lysosomal membranes and to participate in their repair. We have reported previously that VPS13C KO HeLa cells show alterations of lysosome homeostasis, as revealed by an increased number of lysosomes and by activation of the cGAS-STING pathway of innate immunity, possibly reflecting defective and leaky lysosomes. Consistent with these results, immunofluorescence analysis of another cell type, A549 cells, revealed elevated levels of the lysosomal membrane marker LAMP1 in VPS13C KO cells compared to WT cells (Fig. 6A and Fig. S9A). Moreover, fluorescence intensity analysis of cells incubated with LysoView™ 633, a probe whose fluorescence inversely correlates with lysosomal pH, indicated lower intensity in VPS13C KO cells (Fig. 6B and Fig. S9B), suggesting decreased acidification of their lumen.

**Fig. 6.**
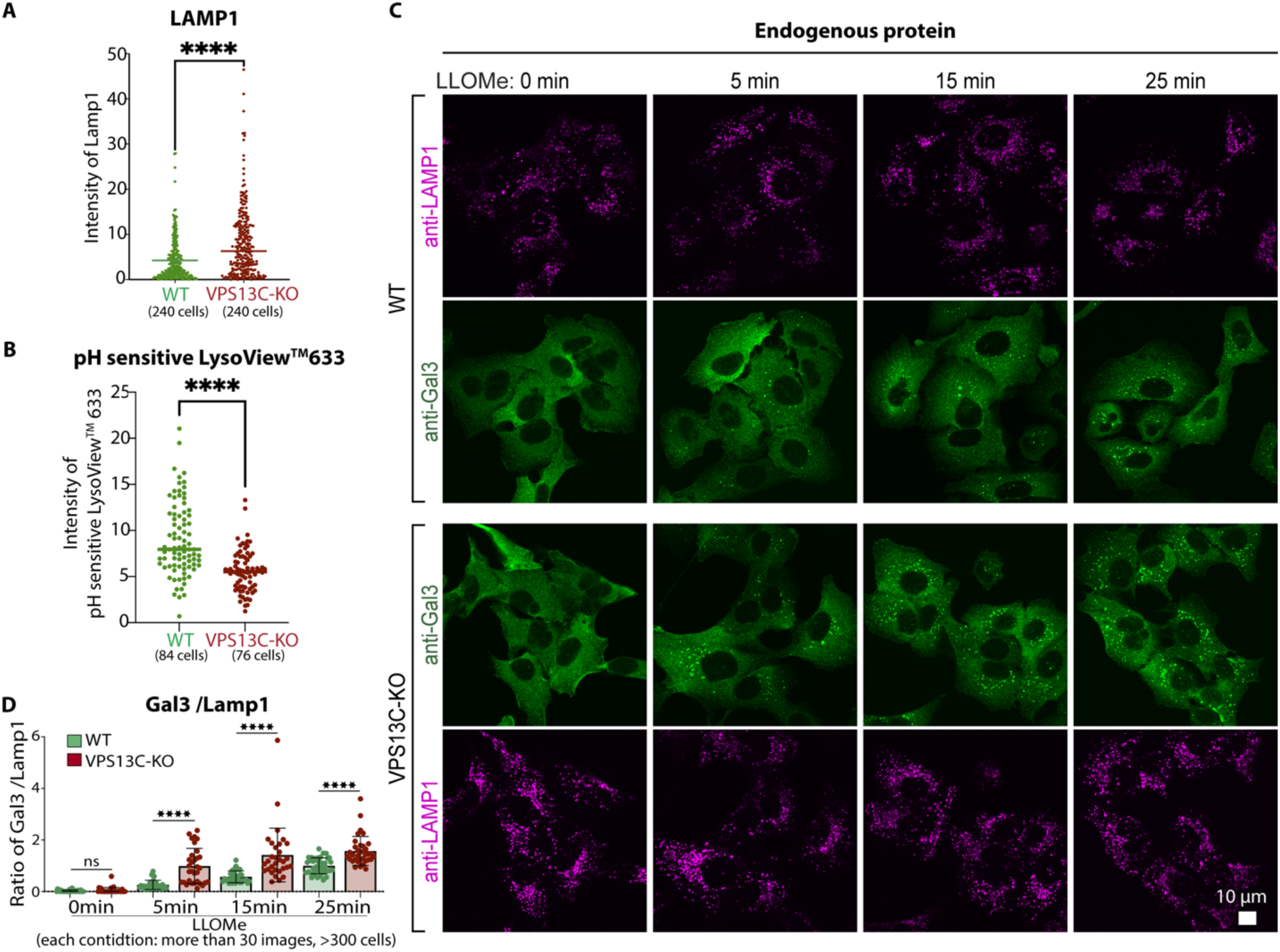
Loss of VPS13C increases the fragility of lysosomes. (A and B) Quantification of the intensity of punctate LAMP1 immunofluorescence (A) or of LysoView633 fluorescence (a PH sensitive lysosome probe) (B) per cell from WT or VPS13C-KO A549 cells. n = 3 biological replicates. Data were compared using two-sided t tests. Error bars represent ±SD. ****, P < 0.0001. (C and D) Fluorescence images of WT and VPS13C-KO A549 cells immunolabeled with antibodies against Gal3 (green) and LAMP1 (magenta) and either untreated or treated with 1 mM LLOMe for the indicated times (C). Quantification of the Gal3 to LAMP1 ratio per image is shown in (D). Bars show the normalized fluorescence relative to WT-LLOMe 25 min. n = 3 biological replicates. Data were compared using two-sided t tests. Error bars represent ±SD. ****, P < 0.0001.

To directly investigate the impact of VPS13C in the response to acute lysosome damage we used the Galectin-3 (Gal3) fluorescence assay. To account for the higher number of lysosomes observed in VPS13C KO cells, all values of Gal3 fluorescence were normalized to LAMP1 immunofluorescence. Under basal conditions, immunofluorescence for endogenous Gal3 did not reveal any accumulation of this protein in either WT or VPS13C KO cells (Fig. 6C, time 0). Upon LLOMe treatment to induce lysosome damage, progressive appearance of Gal3 puncta was observed, and this occurred earlier in VPS13C KO cells compared to WT cells (Fig. 6C and D). Similar results were also observed in HeLa cells by monitoring the fluorescence of transiently expressed Gal3-GFP (Fig. S9C).

Collectively, these findings indicate that the loss of VPS13C increases lysosomal fragility, underscoring its important role in resilience of the lysosomal membranes to damage.

### Relationship between VPS13C and LRRK2 recruitment to damaged lysosomes

LRRK2 can also bind to stressed/damaged lysosomes(*25, 26, 59, 60*). In view of the link of both LRRK2 and VPS13C to PD, we investigated the temporal and mechanistic relationships between VPS13C and LRRK2 recruitment. As various perturbations that induce LRRK2 recruitment and activation do so by triggering the conjugation of ATG8 to single membranes (CASM) process(*37, 61*), we focused on agents that regulate this pathway (Fig. 7A). Our investigations revealed that, like LLOMe, other experimental manipulations that induce CASM, such as chloroquine (a lysosomotropic reagent) or nigericin (an H+/K+ ionophore), which disrupt the acidic luminal pH of lysosomes and thus may have an indirect effect on the lysosomal membrane, can also recruit VPS13C (Fig. 7B and fig. S10A). Recruitment of VPS13C to lysosomes in response to nigericin was also recently reported by a mass spectrometry analysis of purified lysosomes(*62*). However, saliphenylhalamide (SaliP), another agent that increases lysosomal PH and is a well-characterized CASM and LRRK2 activator(*37, 63, 64*), failed to recruit VPS13C (Fig. 7C top) although it induced a loss of LysoView^TM^633 (a pH indicator) fluorescence proving its effectiveness in elevating lysosomal pH (Fig. 7C bottom). SaliP promotes assembly of the V-ATPase responsible for lysosome acidification by inducing the association of its V0 and V1 components through formation of a covalent adduct, but locks it into a pump inactive state(*64, 65*). Furthermore, overexpression of SopF, a Salmonella effector protein which functions as a potent inhibitor of CASM via its property to block the interaction between V-ATPase and ATG16L1(*63, 66*), did not prevent the recruitment of VPS13C by LLOMe treatment (Fig. 7D). These findings demonstrate that while both VPS13C and LRRK2 are recruited to lysosomes in response to their perturbation, VPS13C recruitment in contrast to LRRK2 is not dependent on CASM.

**Fig. 7.**
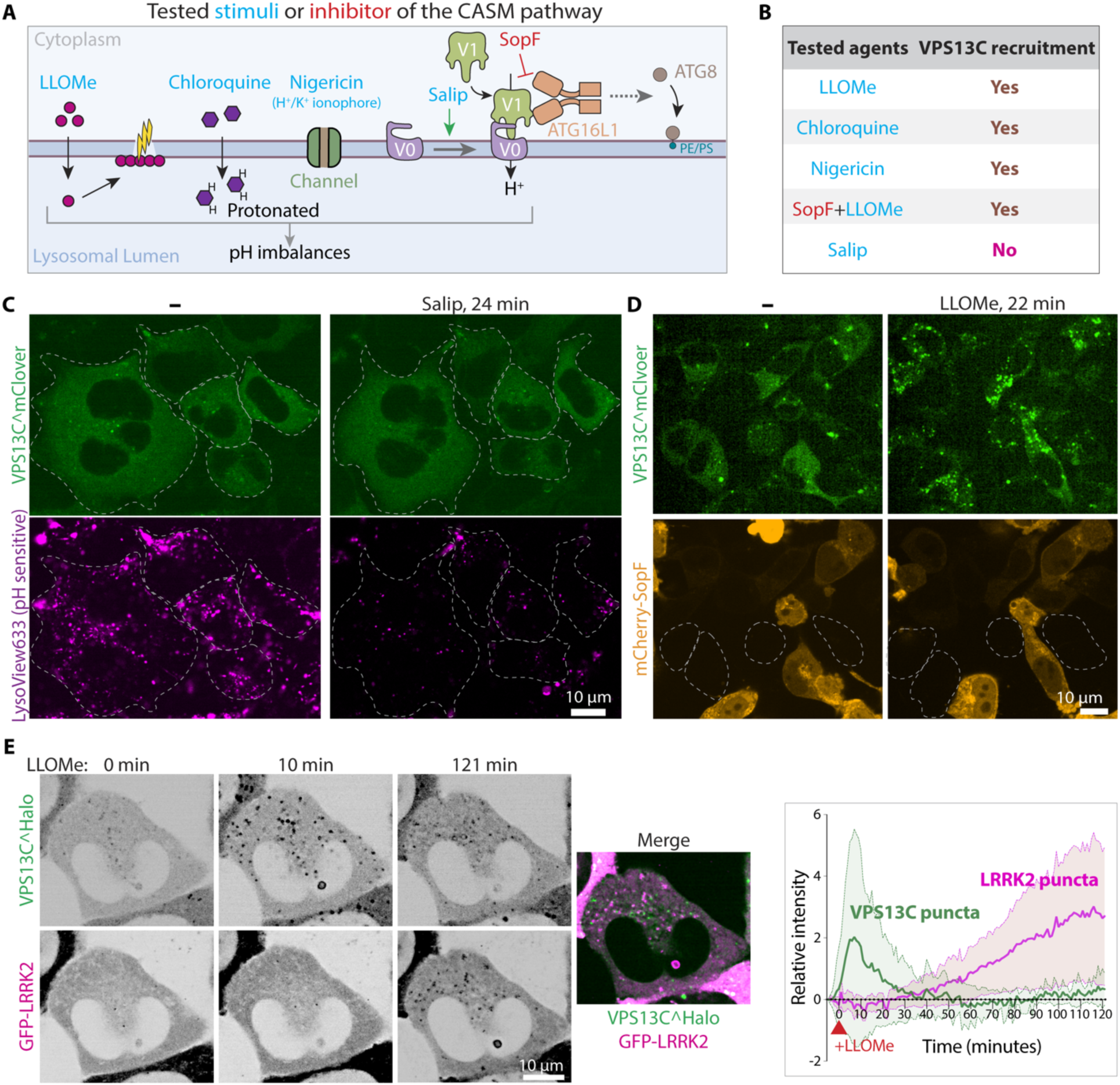
Relationship between VPS13C and LRRK2 recruitment to damaged lysosomes. (A) Schematic drawing illustrating the CASM pathway and experimental manipulations that trigger (blue) or inhibit it (red). (B) Summary of agents tested and their impact on the recruitment of VPS13C to lysosomes. (C) Live fluorescence images of VPS13C^mClover-Flp-In cells labeled with LysoView 633 before and after Salip treatment. (D) Live fluorescence images of VPS13C^mClover-Flp-In cells also co-expressing exogenous mCherry-SopF before and after 1mM LLOMe treatment. Cells not expressing mCherry-SopF are indicated by dashed lines. (E) Time-series of live fluorescence images of Hela cells co-expressing VPS13C^Halo and GFP-LRRK2, showing recruitment of VPS13C and LRRK2 to lysosomes upon 1mM LLOMe treatment. Individual channel images are shown as inverted grays. Quantification of relative punctate fluorescence intensity of VPS13C^Halo or GFP-LRRK2 per cell after LLOMe treatment in a single experiment is shown at the right. >10 cells were analyzed.

We also directly compared the dynamics of VPS13C and LRRK2 recruitment induced by LLOMe. We conducted these experiments in Hela cells, which show robust LLOMe-induced lysosomal recruitment of LRRK2 compared to the VPS13C^mClover-Flp-In cell line used in most of our experiments. Both VPS13C^Halo and GFP-LRRK2 predominantly localized in the cytosol, with VPS13C showing some lysosomal binding under basal conditions. Upon LLOMe treatment, VPS13C was recruited to lysosomes within minutes (Fig. 7E), consistent with observations in VPS13C^mClover-Flp-In cells (Fig. 1B) and RPE1 cells (Fig. 4D, E). Notably, at variance with what we had observed in VPS13C^mClover-Flp-In cells and RPE1 cells, where VPS13C accumulation at endolysosomes persisted for at least 30 minutes after LLOMe addition, VPS13C recruitment in Hela cells was transient, peaking around 10 minutes (Fig. 7E and Fig. S10B). Importantly, recruitment of LRRK2 was much delayed, becoming detectable approximately one hour after addition of LLOMe, when VPS13C had mostly dissociated from lysosomes in these cells (Fig. 7E) and involved a smaller number of lysosomes. A similar sequential recruitment of VPS13C and LRRK2, with fewer lysosomes becoming positive for LRRK2, was also observed in RPE1 cells upon LLOMe treatment (Fig. S10C).

Together, these results demonstrate that although both VPS13C and LRRK2 both respond to lysosome damage, the mechanisms controlling their recruitment differ significantly.

## DISCUSSION

Our study reveals that recruitment of VPS13C is an early response to endolysosome damage, possibly to provide a path for bulk lipid transport from the ER to their membranes. This suggests that lipid transport mediated by this protein via a bridge-like mechanism may play a role in preventing or repairing the rupture of the lysosomal membrane. Via this mechanism, loss of function mutations in VPS13C may contribute to the development of Parkinson’s disease.

These findings are convergent with the recent report that lysosome damage induces the formation/expansion of ER-lysosomes contacts mediated by ER-anchored OSBP and OSBP-related proteins (ORPs) in a process that is dependent on the damage-triggered accumulation of PI4P on the lysosome membrane, the so-called PITT (phosphoinositide-initiated membrane tethering and lipid transport) pathway(*32–34*). OSBP/ORPs are shuttle-like lipid transport proteins that bind the ER protein VAP via an FFAT motif and PI4P-rich Golgi-endosomal membranes via a PH domain(*67, 68*). The PITT pathway, which is activated with a similar time course as the recruitment of VPS13C, is also thought to help provide phospholipids (specifically PtdSer and cholesterol) to the lysosomal membrane from the ER, but in exchange with the counter-transport of PI4P from the lysosome to the ER(*67*), thus not resulting in net lipid transfer to endolysosomes. While we have found that the recruitment to lysosomes of VPS13C is not dependent on PI4P and thus independent from the PITT pathway, the two processes may synergize, as 1) once an ER-lysosome contact has formed, the population of these contacts by other ER-lysosome tethers will be facilitated and 2) the role of OSBP/ORP proteins in regulating the lipid composition of lysosome membranes may be complementary to the expected role of VPS13C in mediating net lipid flow to such membranes.

Activation of the PITT pathway in response to lysosome damage was reported to be followed by the recruitment of ATG2A, another bridge-like lipid transport protein, at ER-lysosome contacts(*32*). The ORP family protein-dependent PtdSer accumulation on the lysosome membrane and/or CASM were proposed to be responsible for such recruitment. However, our results indicate that the recruitment of VPS13C represents a novel response to lysosome damage that occurs independently from the PITT and CASM pathways (*11, 69, 70*).

Other proteins that accumulate with rapid kinetics at the lysosome surface after damage are ESCRT complex components(*27–29*). Such recruitment was proposed to help mediate lysosome membrane repair, as the ESCRT system mediates the formation of spiral filaments that seals membrane holes in a variety of cellular contexts(*31, 71–75*). Thus, the delivery of membrane lipids may function cooperatively with the ESCRT system in membrane resilience or repair. It is also of interest that one of the proposed functions of the single yeast VPS13 protein is to provide phospholipids to endolysosomes for the generation of intraluminal vesicles, an ESCRT-dependent process(*76*). However, whether such a function applies to mammalian VPS13C and, if so, how such a function is related to the recruitment of VPS13 in response to lysosome stress remains to be elucidated.

While VPS13C recruitment requires Rab7, which binds the VAB domain of VPS13C, it also requires an additional signal. We have ruled out Rab7 phosphorylation, PI4P and Ca²⁺ as the signal. We have found instead that such signal is a perturbation of the lysosomal membrane that triggers binding of the ATG2C_VPS13C_ domain, most likely resulting in the release of an autoinhibited conformation of VPS13C that prevents its access to Rab7. Both the VAB domain alone, and VPS13C constructs which contain this domain but lack the ATG2C and PH domains, bind efficiently to endolysosomes also under basal conditions. In contrast, full-length VPS13C and its C-terminal fragment (VPS13C_C-ter_), which contain both the VAB domain and the ATG2C-PH region, do not bind Rab7 in basal conditions. A potential clue about the mechanism that mediates the binding of the ATG2Cdomain of VPS13C to a damaged membrane comes from its known property (and the property of full length VPS13C) to bind lipid droplets via its amphipathic helices (*10*). Many proteins that bind lipid droplets were shown to bind these organelles by recognizing packing defect present in the monolayer that surrounds them(*53, 54*) and this was reported to be the case also for the ATG2C domain of VPS13C(*10*). Bilayer packing defects generated by lysosome damage may be the signal that triggers ATG2C_VPS13C_ recruitment. Supporting this possibility, it was shown that SPG20, another protein that contains amphipathic helices, is also recruited to lysosomes within minutes upon damage of their membranes (*77, 78*). Interestingly, the lysosomal recruitment of SPG20, like that of VPS13C, is independent of PI4P and Ca²⁺ signaling(*77*).

Finally, while both VPS13C and LRRK2 are responsive to lysosomal stress/damage, our study reveals distinct mechanisms and kinetics in their recruitment. VPS13C exhibits rapid recruitment, suggesting an early response in membrane repair/protection, whereas LRRK2 recruitment, which is linked to the activation of CASM, is delayed, pointing to its involvement in late-stage processes, potentially resulting from severe lysosomal damage. Moreover, LRRK2 is recruited to a smaller number of lysosomes. However, a function of both proteins in a response to lysosome damage strengthens evidence(*20–22*) that impairments of such response in vulnerable cell types are critical contributors to the development of PD.

## Acknowledgments

We thank Michael Hanna for advice and discussion and Phyllis Hanson for the kind gift of the IST1 clone. The TBK1/IKKε double knockout MEFs were a kind gift from Shizuo Akira (Department of Host Defense, University of Osaka, Japan). This work was supported in part by grants from the NIH (DA018343, NS36251 to P.D.C.; GM105718 to S.M.F), by the Parkinson’s Foundation (PF-RCE-1946) to P.D.C and S.M.F, by the UK Medical Research Council (grant number MC_UU_00018/1) to D.R.A. and by the Aligning Science Across Parkinson’s grants through the Michael J. Fox Foundation for Parkinson’s Research to P.D.C. and S.M.F (ASAP-000580) and D.R.A. (ASAP-000463). For the purpose of open access, the authors have applied a CC BY public copyright license to all Author Accepted Manuscripts arising from this submission.

## Author contributions

Conceptualization: XW, PDC, SMF, DRA

Investigation: XW, PX, ABD, WHC, SC, BTJ, FT, GT, LS

Supervision: PDC, SMF, DRA

Writing—original draft: XW, PDC

Writing—review & editing: XW, PDC, SMF, DRA

## Competing interests

Authors declare that they have no competing interests.

## Data Availability

The data, code, protocols, and key lab materials used and generated in this study are listed in a Key Resource Table alongside their persistent identifiers at Zenodo DOI:10.5281/zenodo.14846056.

An earlier version of this manuscript was posted to BioRxiv on June 08, 2024 at DOI: https://doi.org/10.1101/2024.06.08.598070

## MATERIALS AND METHODS

### DNA plasmids

A plasmid containing codon-optimized cDNA encoding human VPS13C, with an internal Halo protein after amino acid residue 1914, was generated by and purchased from GenScript Biotech. Plasmids lacking the C-terminal ATG2C and PH domains of VPS13C, designated as VPS13C-Δ(ATG2C-PH)^Halo, as well as those expressing the ATG2C or PH domain of VPS13C (mCherry-ATG2C_VPS13C_ or mCherry-PH), were generated and purchased from Epoch Life Science, Inc. Halo-SspB-PPM1H was constructed as follows: First, the Halo tag was amplified by PCR from the VPS13C^halo plasmid. Subsequently, the stringent starvation protein B (SspB) peptide sequence was added by overlapping PCR, and the construct was ligated into the 3xflag-CMV10 vector using InFusion Cloning, employing the SacI and NotI cut sites. Next, PPM1H was amplified by PCR from the HA-PPM1H plasmid and inserted downstream of SspB using InFusion Cloning with the NotI and BamHI cut sites. The following plasmids were previously generated in our laboratories: VPS13C^mClover3, mCherry-VAB and mCherry-VPS13C_C-ter_ (Kumar et al., 2018); mCherry-VAPB and mCherry-OSBP (Dong et al., 2016); GFP-LRRK2 (Wang et al., 2023); HA-PPM1H (DU62789, MRC Reagents and Services), GFP-LRRK1 (DU30382, MRC Reagents and Services), GFP-LRRK1^K746G^ (DU67083, MRC Reagents and Services) and GFP-LRRK1^D1409A^ (DU67084, MRC Reagents and Services). HA-PPM1H and GFP-LRRK1 constructs were further subcloned into the pmScarlet vector between XhoI and BamHI using InFusion Cloning. The following constructs were obtained from Addgene: mCherry-SopF from L. Knodler (Addgene, #135174); LAMP1-mCherry-iLID from L. Kapitein (Addgene, # 174625); Lamp1-RFP from W. Mothes (Addgene, #1817); mCherry-Gal3 from H. Meyer (Addgene, #85662). IST1-Apple was a kind gift from Phyllis Hanson (University of Michigan School of Medicine, Ann Arbor, MI).

### Antibodies and reagents

#### Antibodies

Anti-Lamp1 [Cell Signaling Technology, 9091; for western blotting (WB), 1:2000], anti-LAMP1 [Abcam, ab25630, for immunofluorescence (IF), 1:100], anti-Galectin3 (R&D Systems, IC1154G, for IF, 1:50), anti-VPS13C (Proteintech, 29844-1-AP; for WB, 1:1000), anti-GM130 (BD Biosciences, 610822; for WB, 1:2000,), anti-PDI (CST, 2446S; for WB, 1:1000), anti-VAPB (Sigma-Aldrich, HPA013144; for WB 1:4000), anti-Rab7 (Cell Signaling Technology, 9367; for WB, 1:1000), mouse monoclonal anti-Rab7A (Sigma Aldrich, R8779; for WB: 1:2000), anti-pSer72 Rab7(Abcam, ab302494; for WB, 1:1000), anti-GAPDH (Proteus, 40-1246; for WB, 1:1000), anti-GFP (Abcam, ab290; for WB, 1:1000), anti-tubulin (Sigma Aldrich, T5168; for WB, 1:2000), anti-mCherry (Abcam, Ab125096; for WB, 1:1000), anti-LRRK1 (MRC Reagents and Services, S405C; for WB: 1 µg/mL), anti-IKKe (Cell Signaling Technology, 3416S; for WB: 1:2000), anti-TBK1 (Cell Signaling Technology, 3504S; for WB, 1:2000), anti-pSer172 TBK1 (Cell Signaling Technology, 5483S; for WB, 1:1000). Secondary antibodies were from LICORbio [IRdye 800CW (926–32213), IRdye 680LT (926–68020); for WB, 1:10,000].

#### Reagents and final concentrations

1mM Leu-Leu methyl ester hydrobromide (LLOMe) (Sigma-Aldrich, L7393-500MG, CAS: 16689-14-8), 200 μM Chloroquine (Sigma-Aldrich, C6628, CAS: 50-63-5), 10 μM Nigericin (Sigma-Aldrich, N7143, CAS: 28643-80-3), 2.5 μM Saliphenylhalamide (Salip) (Omm Scientific), 20nM OSW-1 (MedChem Express, HY-101213, CAS: 145075-81-6), LysoView633 (Biotium, 70058, 1:2000), LysoView640 (Biotium, 70085, 1:2000), 100nM Human RAB7A siRNA (Horizon Biosciences, M-010388-00-0005), 0.1 μg/ml Tetracycline (Thermo Scientific, A39246, CAS: 64-75-5), 200 μM E64d (Cayman Chemical, 13533, CAS: 88321-09-9). Lipofectamine RNAiMAX (Thermo Scientific, 13778030, 1:3), FuGene HD (Promega, E2311, 1:3).

### Generation of tetracycline-inducible Flp-In 293 stable cell line expressing VPS13C^mClover (VPS13C^mClover-Flp-In cell line)

The Flp-In TREx 293 cell line (Invitrogen), which contains a single Flp recombination target (FRT) site in its genome, was maintained in Dulbecco’s modified Eagle’s medium (DMEM), 10% fetal bovine serum (FBS), 200 μg/ml zeocin, and 15 μg/ml blasticidin S. To establish the tetracycline-inducible cell line, pcDNA5/FRT/TO vector containing the cDNA sequence of VPS13C^mClover and pOG44 vector (Invitrogen), which expresses Flp recombinase, were co-transfected into Flp-In TREx 293 cells using FuGene HD (Promega), according to the manufacturer’s recommendations. pcDNA5/FRT vector only was used for transfection as a negative control. At 24 hours post-transfection, cells were washed and fresh media containing Blasticidin but not Zeocin was added. At 48 hours post-transfection, 200 μg/ml hygromycin B (Invitrogen) was added to the culture for selection. The media were changed every 3-4 days until hygromycin-resistant colonies were evident. Expression of the protein was induced by addition of 0.1ug/ml tetracycline in the medium for 24 hours and checked on SDS-PAGE gels followed by Western blots analysis. A detailed method can be found in protocols.io: <u>dx.doi.org/10.17504/protocols.io.36wgqnnnxgk5/v1</u>

### Generation of a VPS13C KO A549 cell line

A549 cells were transfected with 1.5 μg of PX459 plasmid (plasmid #62988; Addgene) containing a sgRNA(*1*) against VPS13C using Lipofectamine 2000 (Thermo Fisher Scientific). At 24 h after transfection, cells were selected in complete DMEM containing 1 μg/ml puromycin. At 48 and 72 h after transfection, medium was replaced with fresh puromycin-containing medium. After 3 days of puromycin selection, single clones were obtained using serial dilution and then screened by western blotting (WB). Cells lacking VPS13C by WB were selected.

### Generation and culture of mouse embryonic fibroblasts

Wild type and homozygous LRRK1 knockout MEFs(*2*) were isolated from littermate matched mouse embryos at day E12.5 resulting from crosses between heterozygous LRRK1 KO/WT mice using the protocol described in: dx.doi.org/10.17504/protocols.io.eq2ly713qlx9/v1.

MEFs generated from double-knockout mice that do not express TBK1 and IKKɛ were a kind gift from Shizuo Akira (Department of Host Defense, University of Osaka, Japan) and were described previously(*3*). Genotypes were verified via allelic sequencing and immunoblotting analysis. Cells were cultured in DMEM containing 10% (v/v) FBS, 2 mM L-glutamine, penicillin-streptomycin 100 U/mL, 1 mM sodium pyruvate, and 1× non-essential amino acid solution (Life Technologies, Gibco). Cells were regularly tested for Mycoplasma PCR products using a Lonza Mycoplasma kit.

### Cell culture and transfection

VPS13C^mClover-Flp-In cells and Hela cells were cultured at 37°C with 5% CO2 in DMEM medium supplemented with 10% FBS. RPE1 cells were cultured at 37°C with 5% CO2 in DMEM/F12 medium supplemented with 10% FBS. For live-cell imaging experiments, cells were seeded onto glass-bottomed dishes (MatTek). For biochemical experiments, cells were plated in 6cm-diameter dishes. Following 24 hr incubation, cells were transfected using FuGene HD (Promega) according to manufacturer’s recommendations. Concurrently, 0.1ug/ml tetracycline (working concentration) was added to the medium to induce VPS13C^mClover expression and cells were imaged 20-24 hr later. Transfection of siRNA was carried out using Lipofectamine RNAiMax (Thermo Fisher Scientific) with 100 nM siRNA pool per the manufacturer’s recommendations and cells were either imaged or collected for Western blot analysis 48 hr later.

### Live-cell imaging

Growth media were changed with live-cell imaging solution (Life Technologies) shortly before imaging. Imaging was performed at 37°C in 5% CO2 with a Nikon Ti2-E inverted microscope equipped with Spinning Disk Super Resolution by Optical Pixel Reassignment Microscope (Yokogawa CSU-W1 SoRa, Nikon) and Microlens-enhanced Nipkow Disk with pinholes and 60x SR Plan Apo IR oil immersion objective. For LysoView experiments, cells were incubated in complete DMEM for 30 min, washed twice with fresh media, and then imaged in live-cell imaging solution. For LLOMe experiments, a 2x stock solution was prepared in the imaging solution and added to the dish during imaging.

For the light-dependent recruitment of PPM1H to lysosomes, the same Yokogawa CSU-W1 SoRa microscope was used. Recruitment was achieved with 200-ms pulses of the 488-nm laser. Imaging was performed at 37°C in 5% CO2, with a 60x SR Plan Apo IR oil immersion objective.

A detailed description of cell culture, transfection, and imaging can be found in protocols.io: dx.doi.org/10.17504/protocols.io.eq2lyp55mlx9/v1

### Immunofluorescence

WT or VPS13C-KO A549 cells grown on glass coverslips and fixed with 4% PFA in PBS (Gibco, 14190144) for 20 min at room temperature, washed 3x with PBS, permeabilized with 1xTris Buffered Saline with Tween buffer (Santa Cruz Biotechnology, sc-281695) for 30 min at room temperature, and blocked using filtered PBS containing 3% (wt/vol) BSA for 1 h at room temperature. Coverslips were then incubated with antibodies, LAMP1 (H4A3) (Abcam, ab25630,1:100) and Galectin-3 Alexa Fluor 488-conjugated (R&D Systems, IC1154G, 1:50) at 4°C overnight, followed by 3× washes in PBS. Secondary antibodies (1:1,000, Alexa Fluor 546, Invitrogen) were incubated in PBS containing 3% BSA for 1 h at room temperature in the dark and washed by 3× PBS. Finally, coverslips were mounted onto slides.

### Generation of Superparamagnetic Iron Oxide Nanoparticles (SPIONs)

SPIONS were generated based on an established protocol(*1, 4, 5*). Briefly, 10 mL of 1.2M FeCl2 (Sigma-Aldrich, 220299) and 10 mL of 1.8M FeCl3 (Sigma-Aldrich, 157740) were combined slowly by stirring. Then, 10 mL of 30% NH4OH (Sigma-Aldrich, 320145) was slowly added while stirring for 5 minutes. The resulting particles were then washed with 100 mL of water three times. The particles were resuspended in 80 mL of 0.3M HCl (J.T. Baker, 9535) and stirred for 30 minutes.

Then, 4g of dextran (Sigma-Aldrich, D1662) was added and stirred for 30 minutes. The particles were transferred into dialysis tubing (Thermo Fisher, 68100) and dialyzed with ddH2O for at least 2 days with multiple water changes. The particles were centrifuged at 26,900g for 20 minutes to remove large aggregates and stored at 4°C until further use. A detailed method can be found in protocols.io: <u>dx.doi.org/10.17504/protocols.io.eq2lyn69pvx9/v1</u>

### Isolation of lysosomes using SPIONs

Cells were cultured in 15-cm dishes until reaching 90-95% confluency, followed by removal of media and a single wash with PBS (Thermo Fisher, 10010023). Subsequently, 20 mL of fresh DMEM media (supplemented with FBS and Pen/Strep) containing 10% SPIONs particles and 10 mM HEPES pH 7.4 (Thermo Fisher, 15630080) was added, and cells were then incubated at 37°C for 24 hours. After the incubation, the media containing particles was removed, and cells were washed once with PBS. Cells were trypsinized (Thermo Fisher, 25200056), re-plated evenly into four 10-cm culture dishes and incubated for 24-hour at 37°C. Upon removal of media, cells were washed three times with PBS. Cells were collected in PBS by scraping on ice, followed by centrifugation at 1,000 rpm for 5 minutes at 4°C. The cell pellet was then resuspended in 1 mL of homogenization buffer (HB) per dish, containing 5 mM Tris (pH 7.4, RPI, T60040), 250 mM sucrose (Sigma, S0389), 1 mM EGTA (Sigma-Aldrich, E4378), and phosphatase/protease inhibitors (PhosSTOP Roche 4906837001 / cOmplete mini EDTA Free Roche 11836170001). Whole cell lysate (WCL) was generated using a dounce homogenizer (DWK Life Sciences, 357538), followed by centrifugation of the WCL at 800g for 10 minutes at 4°C to collect the supernatant (WCL). The WCL was loaded onto a LS column (Miltenyl Biotec, 130042401), and the flow-through was collected and reapplied to the same column. After washing the column once with 3 mL of HB, the column was removed from the magnetic stand, and the bound fraction was eluted using 2.5 mL of HB. The bound fraction was then centrifuged at 55,000 rpm for 1 hour at 4°C to generate the lysosomal pellet, which was further resuspended in 50 uL of HB and stored at −20°C until further processing. A detailed method can be found in protocols.io <u>dx.doi.org/10.17504/protocols.io.bp2l61dr1vqe/v1.</u>

### Western blotting (WB)

Cultured cells were lysed on ice through repeated pipetting in 2% SDS supplemented with Protease Inhibitor Cocktail (Roche) and PhosStop phosphatase inhibitor (Roche). Cell lysates were further sonicated for 30 seconds per sample to break DNA and eliminate viscosity of the samples. Total protein was then measured by Pierce BCA assay (Thermo Fisher Scientific). Samples were prepared for WB at equal protein concentrations, mixed with SDS loading buffer [final: Bromophenol blue (0.05%), DTT (dithiothreitol; 0.1 M), Glycerol (10%), SDS (sodium dodecyl sulfate; 2%), Tris-Cl (0.05 M, pH 6.8)] and exposed to 95°C for 5 min. Proteins were separated on Mini PROTEAN TGX 4–20% Tris-glycine gels (Bio-Rad) before transfer to nitrocellulose membranes at 4°C for 2 h at 100 V in transfer buffer containing 25 mM Tris, 192 mM glycine, and 20% methanol in milliQ purified water. Transferred membranes were “blocked” in 5% milk in Tris-buffered Saline (TBS) containing 0.1% Tween-20 (TBST) for 1 h. Membranes were then incubated with primary antibodies in 3% BSA in PBST overnight at 4°C. The next day, membranes were washed 3 times in TBST and then incubated with secondary antibodies conjugated to IRdye 800CW or IRdye 680LT (Licor, 1:10,000) in 5% milk in TBST at RT for 1 h, washed 3 times in TBST, and then imaged using a Licor Odyssey Infrared Imager. For the experiments shown in Figure S3, cells were lysed, and whole cell lysates analyzed by immunoblotting, as described in <u>dx.doi.org/10.17504/protocols.io.ewov14znkvr2/v2</u>.

### Image analysis and statistics

Fluorescence images presented in this study were processed with Fiji software. Quantification of fluorescence puncta (endolysosomes) or diffuse cytosolic fluorescence was performed as follows. First, the images were converted to 8-bit files and separated into individual channels. Subsequently, an automatic alignment of cells was conducted using the StackReg-Translation plugin in Fiji. Next, outlines delineating individual cells were manually drawn and adjustments of image thresholds were manually performed in each channel to ensure consistency across all images within the same channel. To isolate the relative puncta signal, the threshold was adjusted to eliminate all cytosolic signals by elevating the minimum value. Conversely, for the relative cytosolic signal, the threshold was adjusted to remove all puncta signals by reducing the maximum value. The average intensity of each cell over time was calculated using the Time Series Analyzer V3 plugin in Fiji. To obtain the final relative intensity over time, the values from each cell were normalized by subtracting the initial values, thus setting the first value as zero.For quantifying the puncta intensity ratio between the Galectin3 and LAMP1 fluorescent channels, we first needed to differentiate the true fluorescent signal from noise and background (Cytosol).

For quantification of the Gal3 to LAMP1 fluorescence ratios, we used the “Trainable Weka Segmentation” plugin in FIJI and user interaction to train a classifier for each channel. Three datasets were used for training the classifiers, one for each channel. They were then applied to all datasets to mask the Galectin3 and LAMP1 positive regions in the corresponding channel. Within those masked regions, measurements of “Integrated density” using FIJI’s “Measure” tool were performed for each fluorescent channel, after which a ratio between these two measurements were obtained for each dataset using a script available for download at https://github.com/linshaova/xinbo-VPS13C.git.

Statistical analysis was performed with GraphPad Prism 8 software (v8.0.1, http://www.graphpad.com/, RRID: SCR_002798) software.

---

**Fig. S1.**
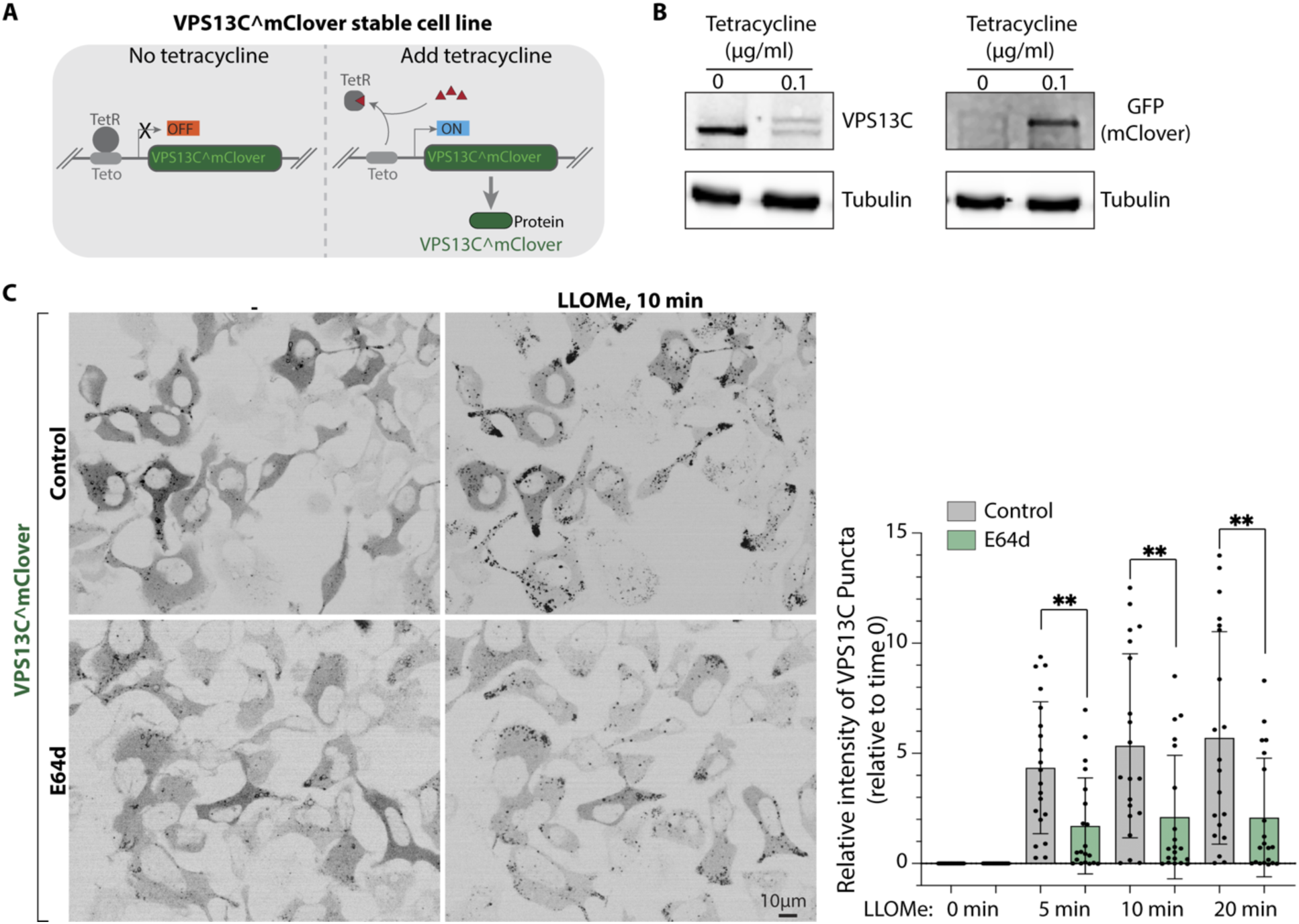
Validation of the VPS13C^mClover-Flp-In cell line under the control of tetracycline and effect of the cathepsin C inhibitor E64D on VPS13C recruitment to endolysosomes. (A) Schematic drawing showing the experimental system used for the inducible stable expression of VPS13C^mClover. (B) Western blots for the indicated proteins of whole cell lysates from VPS13C^mClover-Flp-In cells under the control of tetracycline, with or without tetracycline (0.1 μg/ml) treatment for 24 hours. Tubulin was used as a loading control. (C) Live fluorescence images of VPS13C^mClover-Flp-In cells before and after a 10 min exposure to 1mM LLOMe with and without the additional presence of the cathepsin inhibitor E64d (200 μM). Quantification of the intensity of the VPS13C^mClover punctate fluorescence per cell from a representative experiment is shown on the right. Graph shows the normalized fluorescence relative to time 0. Data were compared using two-sided t tests. Error bars represent ±SD, 20 cells were analyzed. **, P < 0.01.

**Fig. S2.**
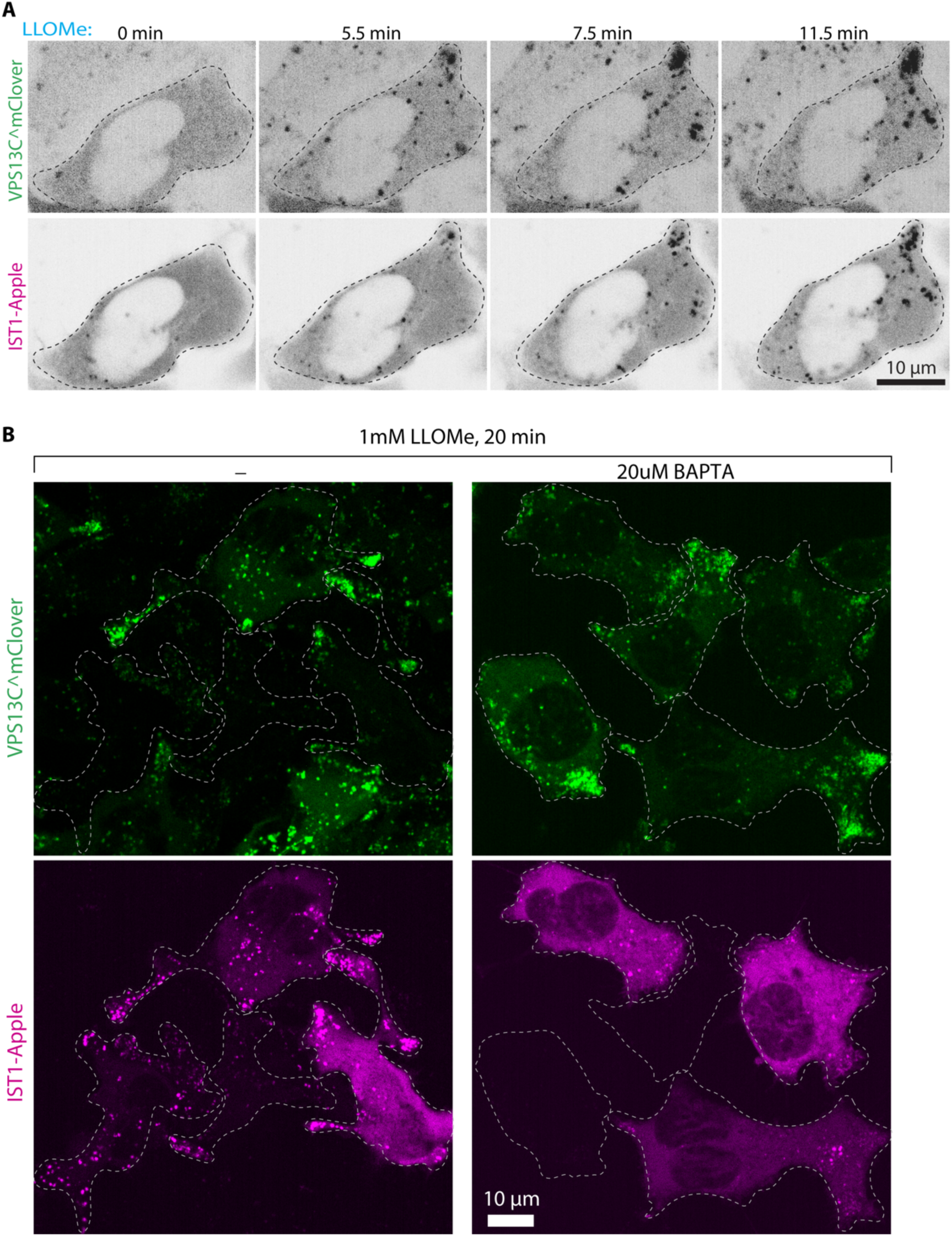
LLOMe-induced lysosomal recruitment of VPS13C precedes of ESCRT-III. (A) Time-series of live fluorescence images of VPS13C^mClover-Flp-In cells also co-expressing IST1-Apple, upon 1mM LLOMe treatment. Note the delayed recruitment of IST1 to lysosomes relative to VPS13C. Individual channel images are shown as inverted grays. (B) Live fluorescence images of VPS13C^mClover-Flp-In cells also co-expressing IST1-Apple, 20 minutes after addition of 1mM LLOMe. Cells were pre-incubated for 1hr with or without the calcium chelator BAPTA (20 μM). Recruitment of IST1, but not of VPS13C, is inhibited by BAPTA.

**Fig. S3.**
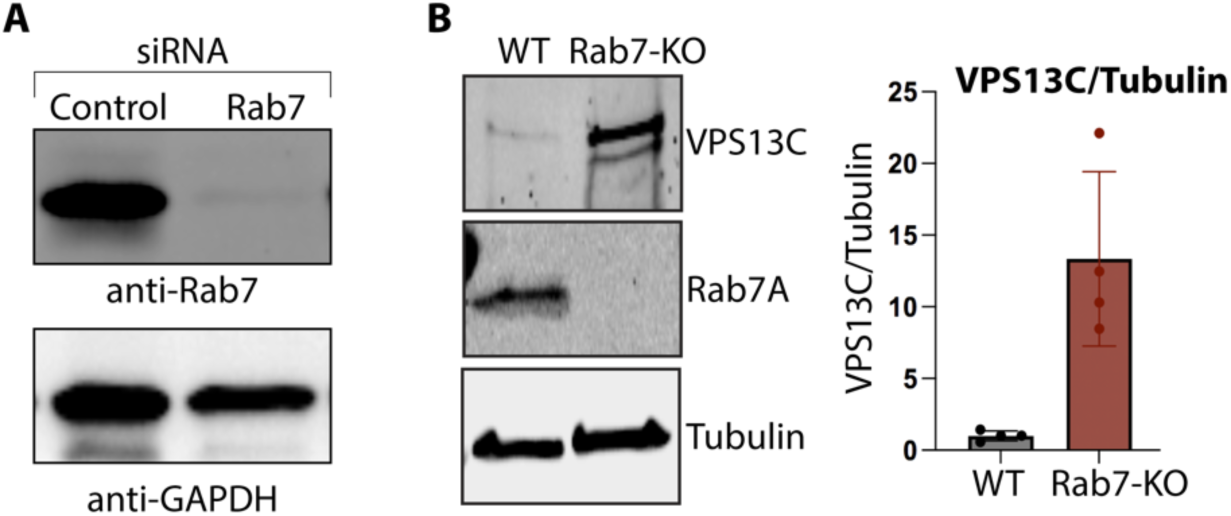
Validation of the Rab7 knockdown or knockout cells. (A) Anti-Rab7 western blot of whole cell lysates from control or Rab7 knockdown VPS13C^mClover-Flp-In cells. GAPDH as a loading control. (B) Anti-Rab7 or anti VPS13C western blots of whole cell lysates from WT or Rab7 knockout Hela cells. Tubulin as a loading control. n = 4 biological replicates.

**Fig. S4.**
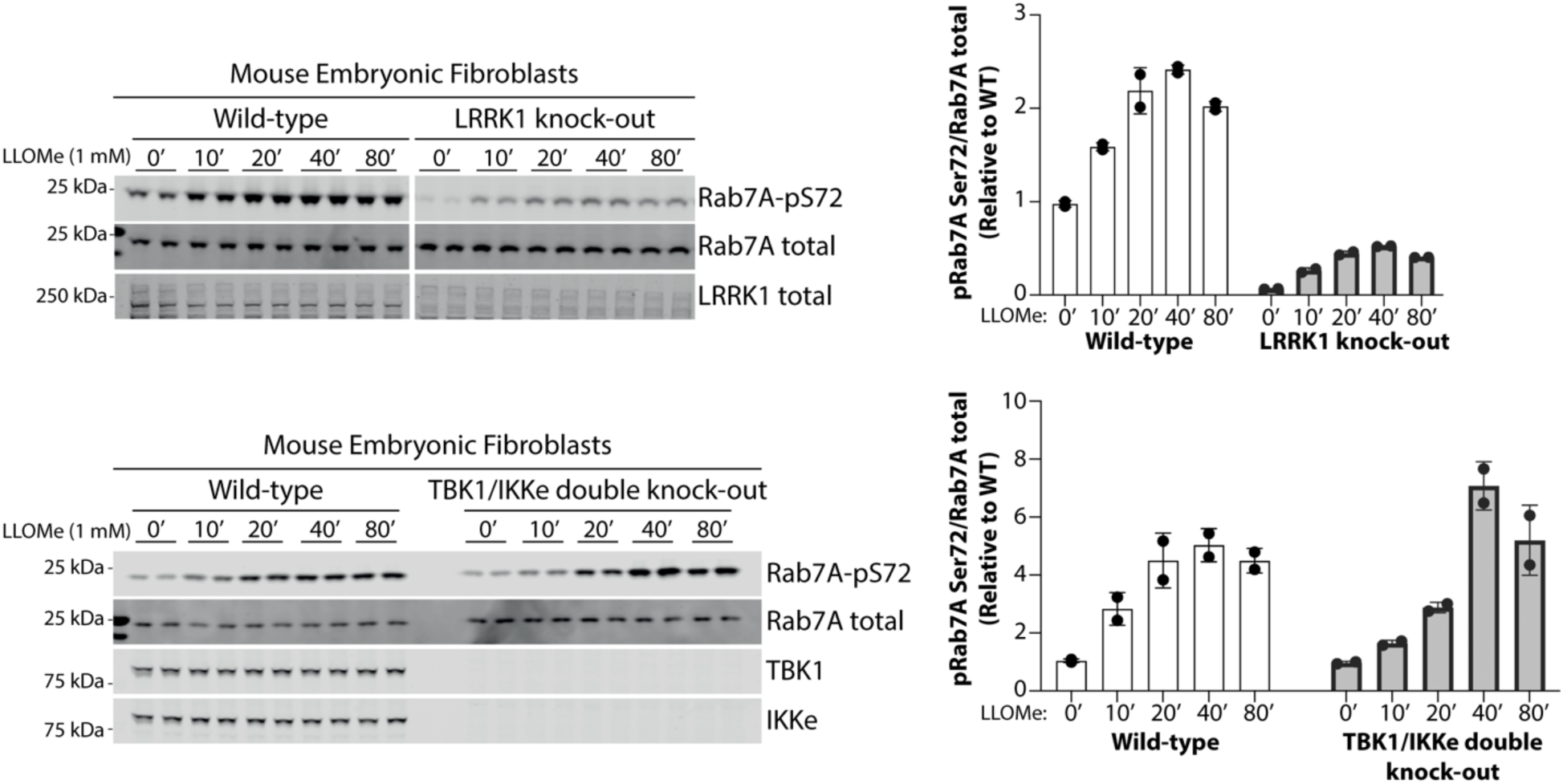
Effect of the KO of LRRK1 and TBK1 on LLOMe-induced Rab7 Ser72 phosphorylation in mouse embryonic fibroblasts (MEFs) Western blot analysis for the indicated proteins in whole cell lysates from the WT, LRRK1 KO or TBK1/IKKe DKO mouse embryonic fibroblasts treated with 1mM LLOMe for the indicated times. Quantification of the data is shown on the right.

**Fig. S5.**
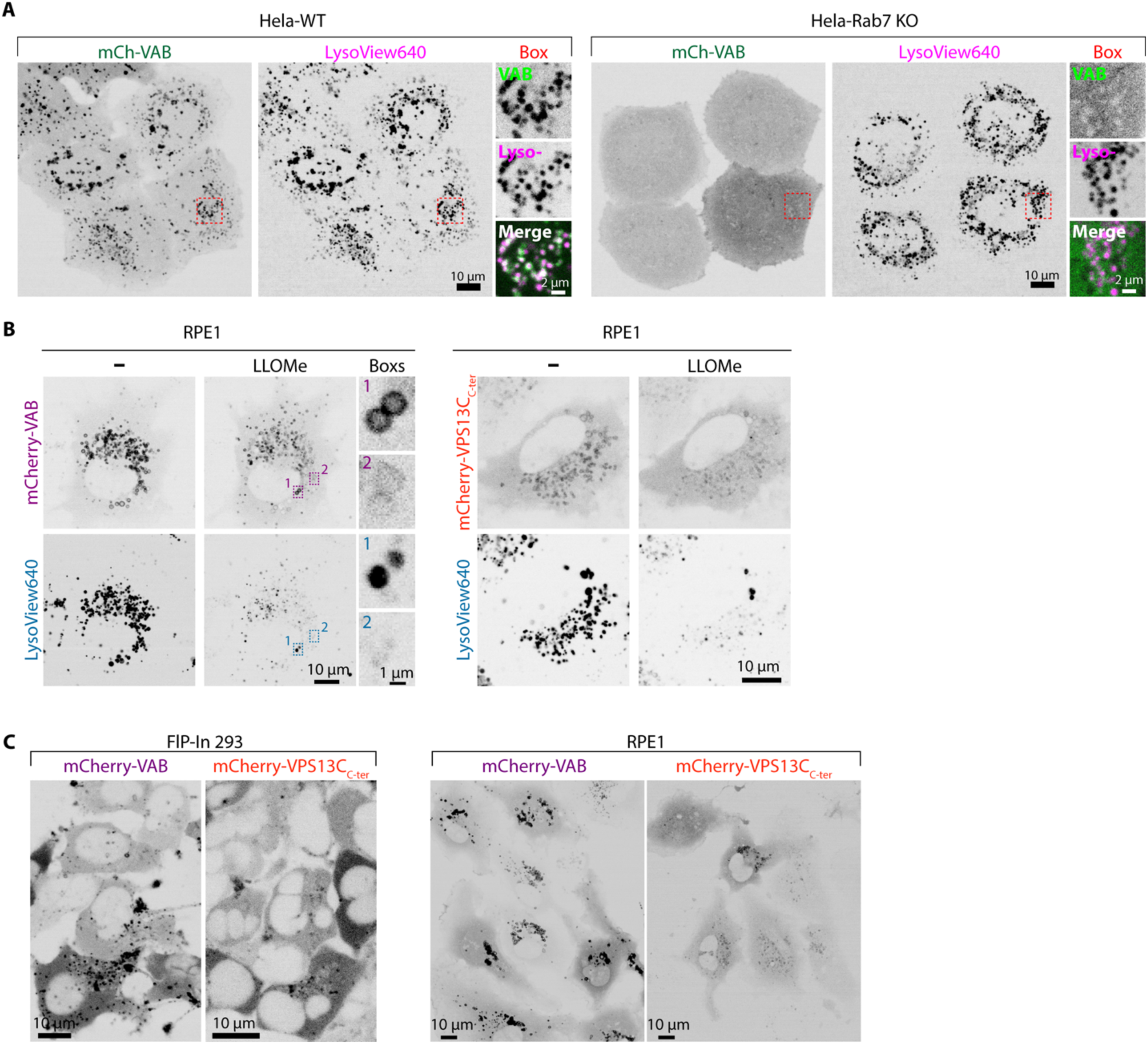
Differential lysosome binding of the VAB domain alone and of the entire C-terminal fragment of VPS13C (VPS13C_C-ter_) (A) Live fluorescence images of WT or Rab7 KO Hela cells expressing mCherry-VAB and labeled with LysoView 640 under basal conditions. Boxed regions are shown on the right. (B) Live fluorescence images of RPE1 cells expressing exogenous mCherry-VAB (left) or mCherry-VPS13C_C-ter_ (right) before and after 1mM LLOMe treatment. (C) Live fluorescence images show localizations of mCherry-VAB or mCherry-VPS13C_C-ter_ in VPS13C^mClover-Flp-In cells (left) or in RPE1 cells (right) in the absence of LLOMe treatment. Note the more prominent lysosome localization of the VAB domain relative to VPS13C_C-ter_.

**Fig. S6.**
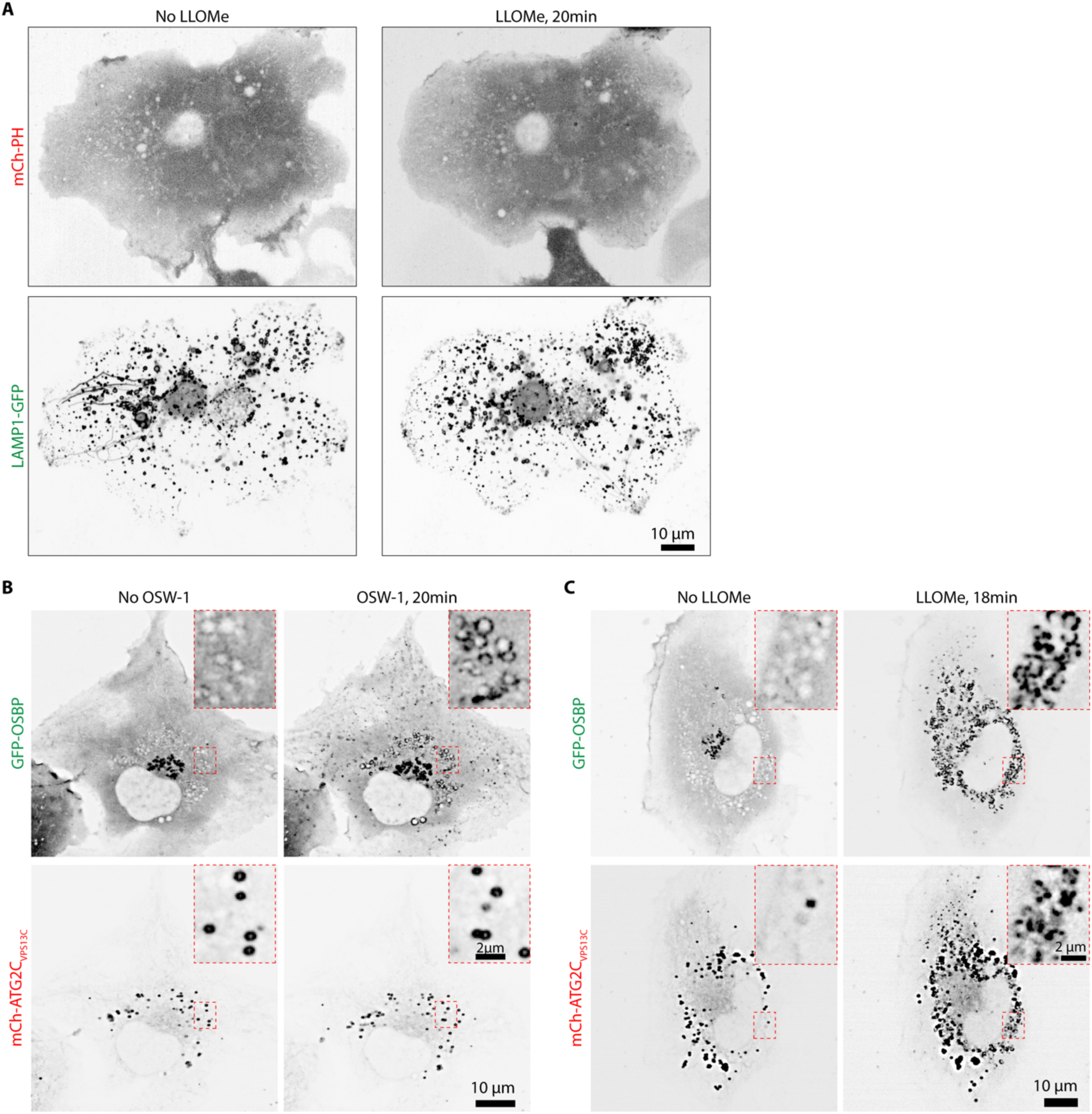
Impact of either LLOMe or OSW1 on the localization of the ATG2C domain or the PH domain of VPS13C. (A) Live fluorescence images of RPE11 cells expressing mCherry-PH and LAMP1-GFP before and after 1mM LLOMe addition. (B and C) Live fluorescence images of RPE1 cells expressing mCherry-ATG2C_VPS13C_ and GFP-OSBP before and after addition of 20nM OSW1 (B) or 1mM LLOMe (C). Black dots visible before LLOMe are lipid droplets that move slightly in position after LLOMe. Individual channel images are shown as inverted grays.

**Fig. S7.**
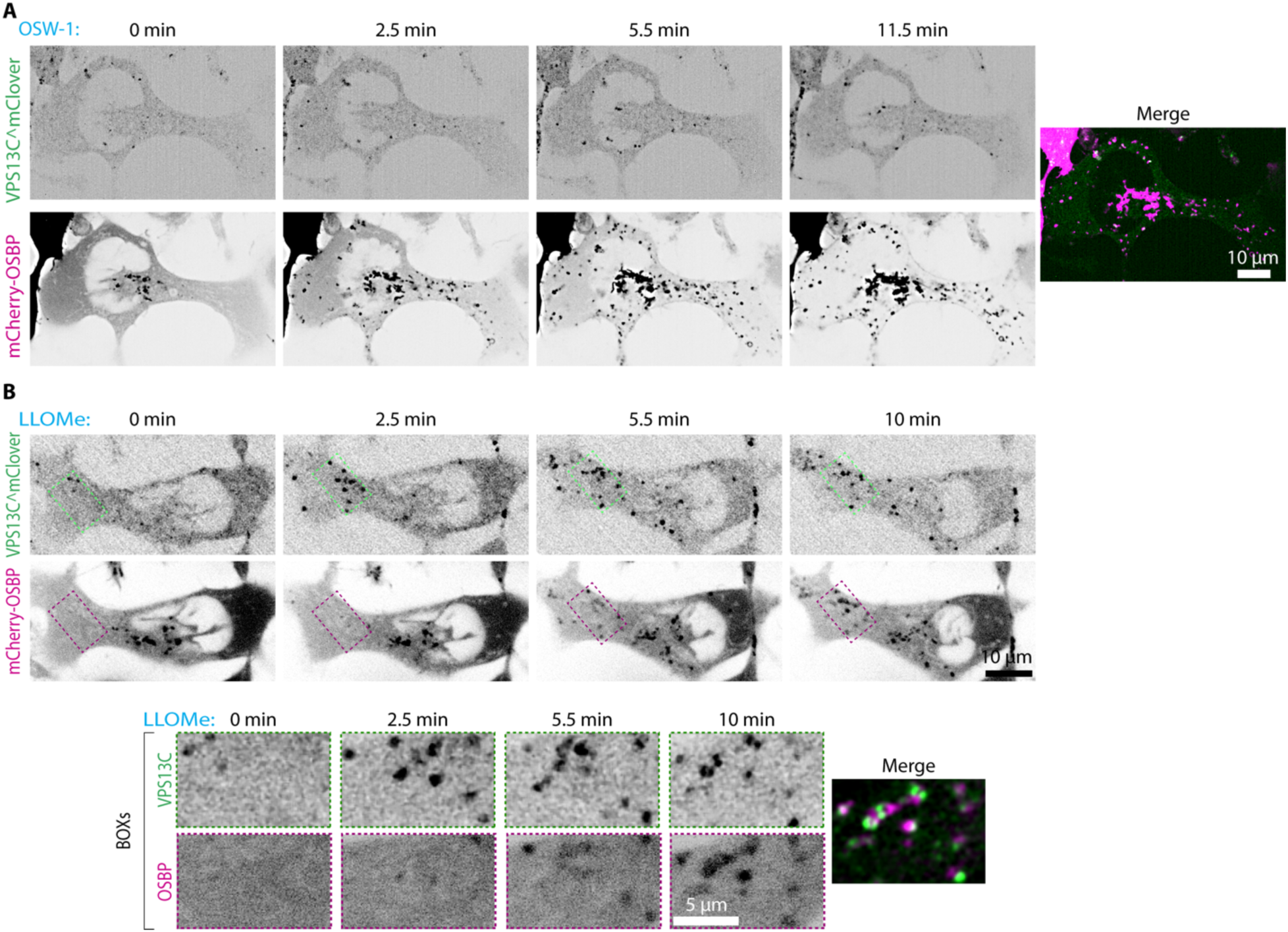
Impact of either LLOMe or OSW1 on the localization of VPS13C or OSBP. (A) Time-series of live fluorescence images of VPS13C^mClover-Flp-In cells also co-expressing exogenous mCherry-OSBP, upon 20nM OSW-1 treatment to induce PI4P accumulation on endolysosomes. Individual channel images are shown as inverted grays. Note that the recruitment of OSBP is not accompanied by the recruitment of VPS13C. (B) Time-series of live fluorescence images of VPS13C^mClover-Flp-In cells also co-expressing exogenous mCherry-OSBP, upon 1mM LLOMe treatment. Note the delayed recruitment of OSBP to lysosomes relative to VPS13C. Individual channel images are shown as inverted grays. Boxed regions are shown at the bottom. Individual channel images are shown as inverted grays.

**Fig. S8.**
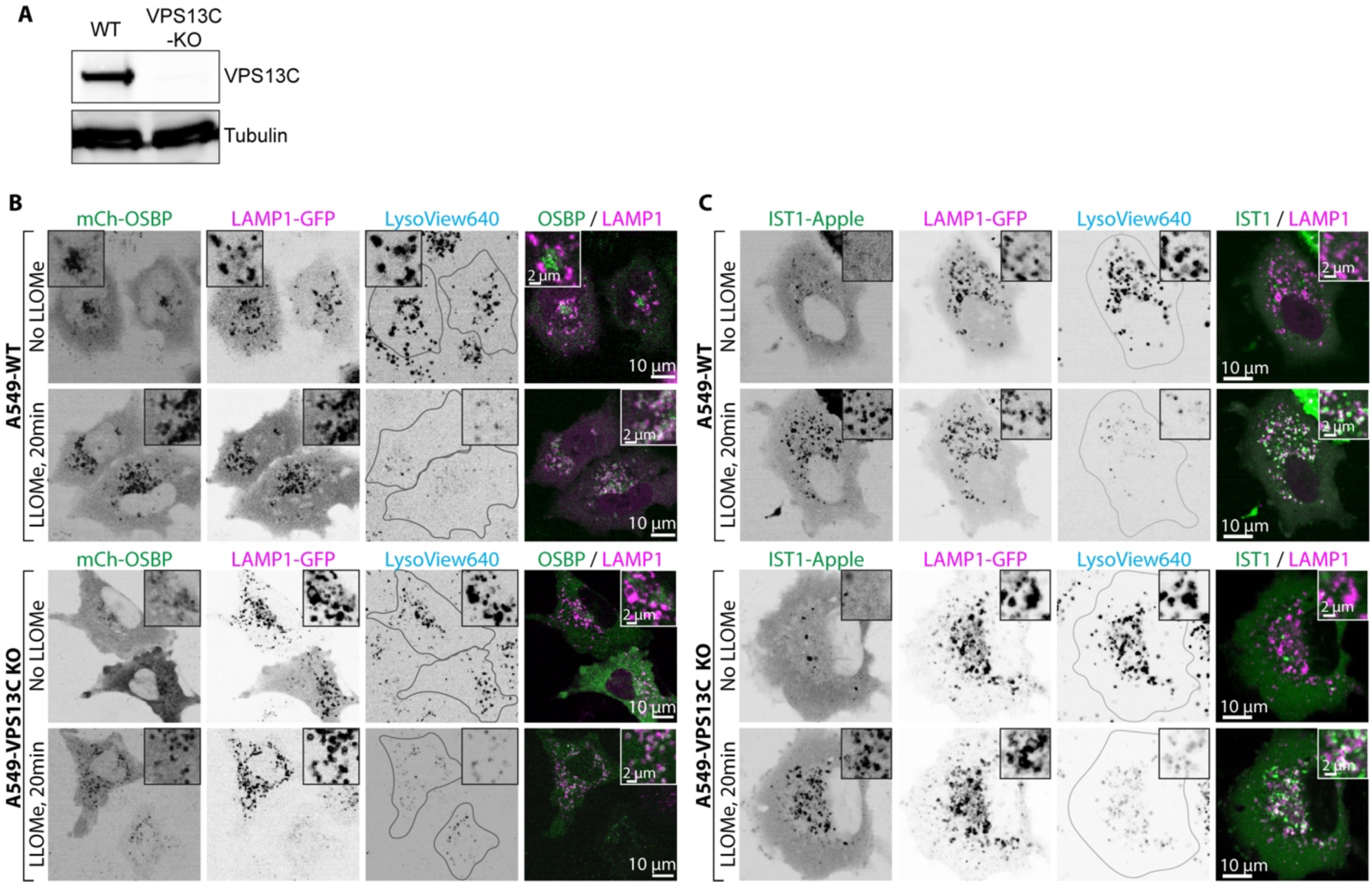
Depletion of VPS13C does not affect the lysosomal recruitment of OSBP and IST1. (A) Anti-VPS13C western blot of whole cell lysates from WT or VPS13C knockout A549 cells. Tubulin was used as a loading control. (B and C) Live fluorescence images of the WT and VPS13C-KO A549 cells expressing mCherry-OSBP and LAMP1-GFP (B) or IST1-Apple and LAMP1-GFP (C) and stained with LysoView640 before and after 1mM LLOMe treatment. Individual channel images are shown as inverted grays.

**Fig. S9.**
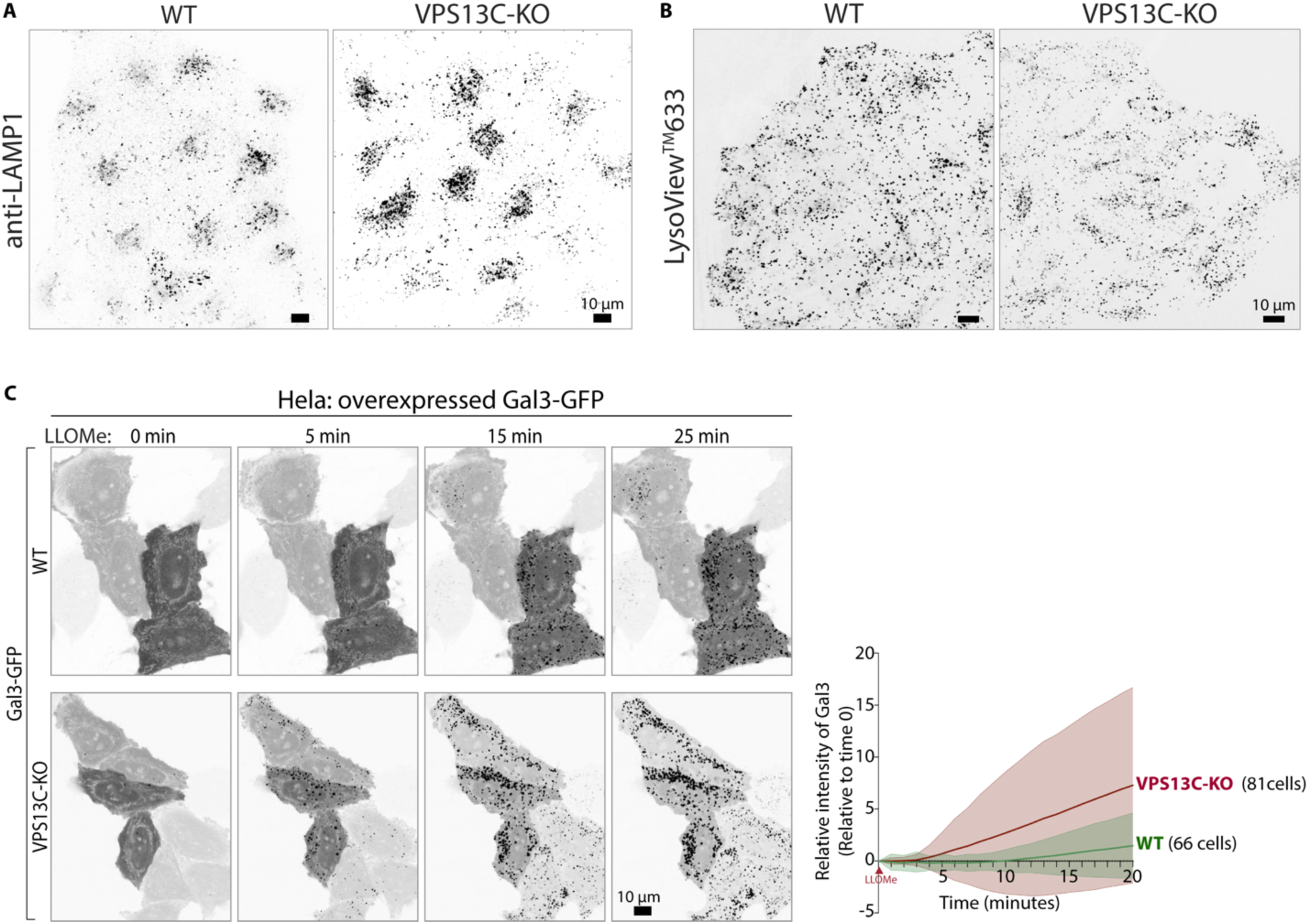
Depletion of VPS13C causes disruption of lysosome homeostasis. (A) Anti-LAMP1 immunofluorescence of WT or VPS13C-KO A549 cells. (B) Live fluorescence images of WT or VPS13C-KO A549 cells incubated with the pH sensitive LysoView633 showing lower fluorescence (higher pH) of the KO cells. (C) Time-series of live fluorescence images of WT or VPS13C-KO Hela cells expressing Gal3-GFP showing recruitment of Gal3 to damaged lysosomes upon 1mM LLOMe treatment. Quantification of the intensity of the Gal3-GFP punctate fluorescence per cell after addition of 1mM LLOMe is shown on the right. n = 3 biological replicates. Individual channel images are shown as inverted grays.

**Fig. S10.**
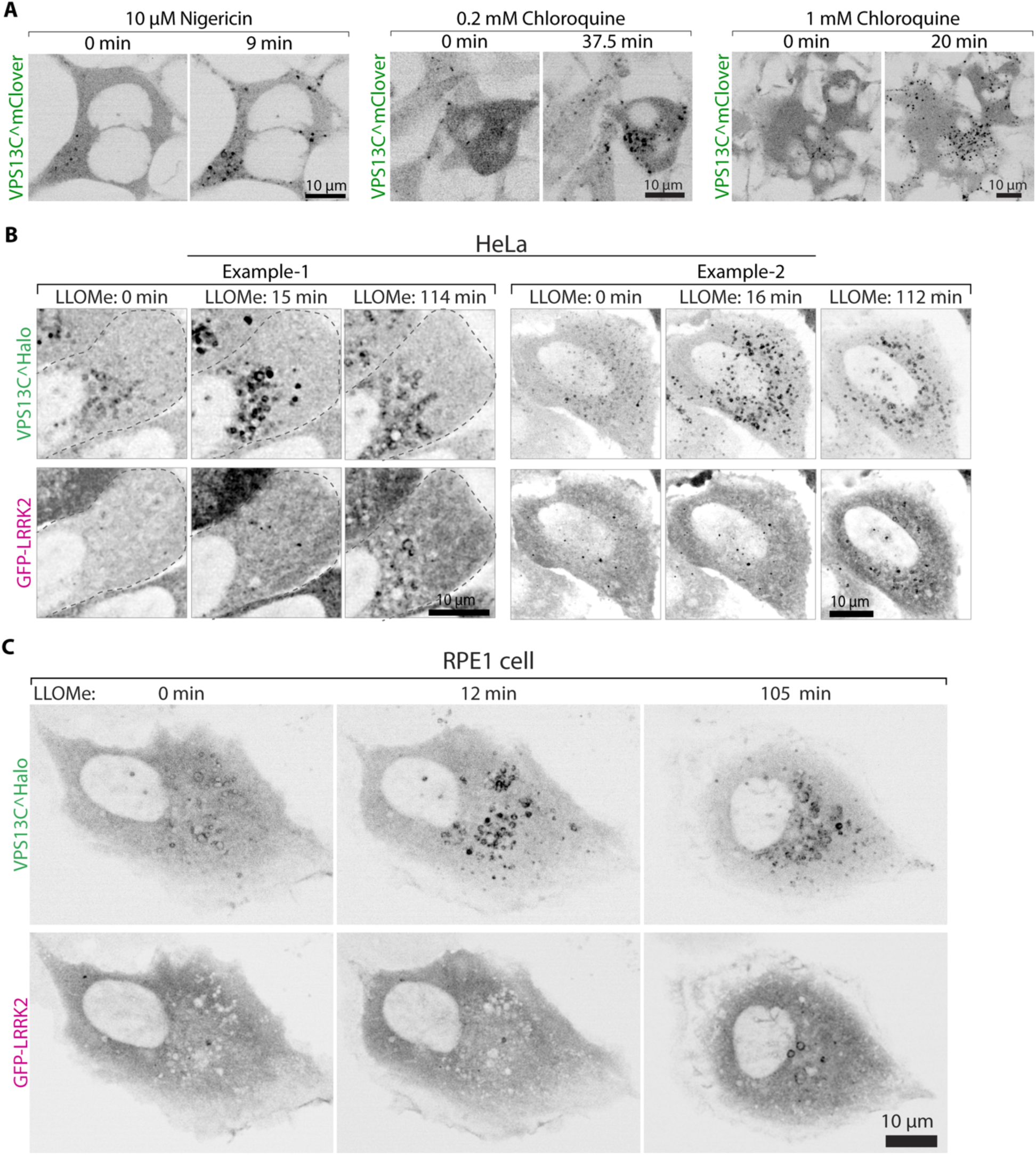
Effect of various agents on lysosomal recruitment of VPS13C or LRRK2. (A) Live fluorescence images of VPS13C^mClover-Flp-In cells before and after addition of nigericin or chloroquine, showing lysosomal recruitment of VPS13C. (B) Time-series of live fluorescence images of Hela cells co-expressing VPS13C^Halo and GFP-LRRK2, showing recruitment of VPS13C and LRRK2 to lysosomes upon 1mM LLOMe treatment. Individual channel images are shown as inverted grays. (C) Time-series of live fluorescence images of a RPE1 cell co-expressing VPS13C^Halo and GFP-LRRK2 showing recruitment of VPS13C and LRRK2 to damaged lysosomes upon 1mM LLOMe treatment. LRRK2 is recruited only to a small subset of lysosomes and with a much slower time course than VPS13C. Individual channel images are shown as inverted grays.

## Key Resource Table

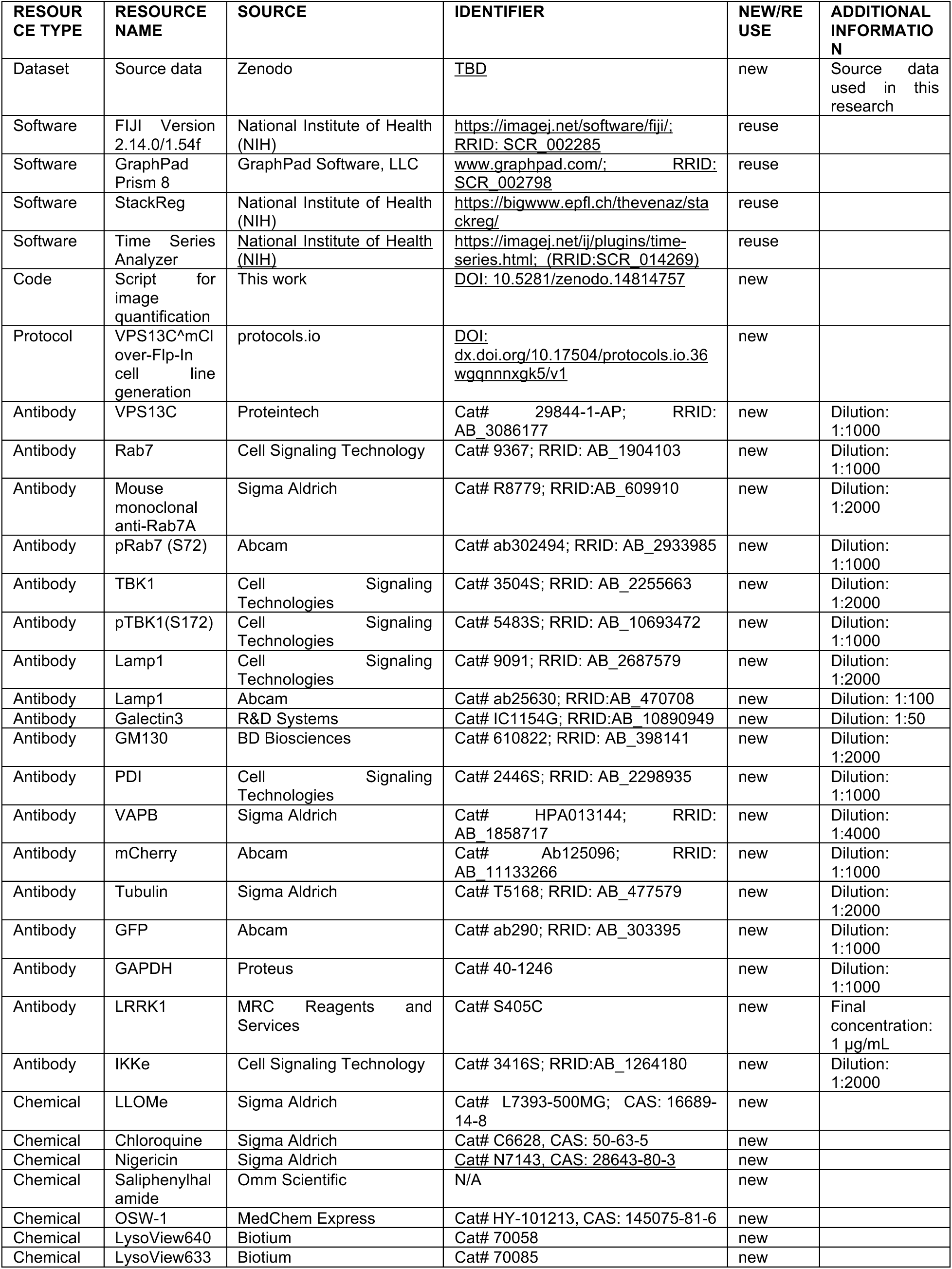
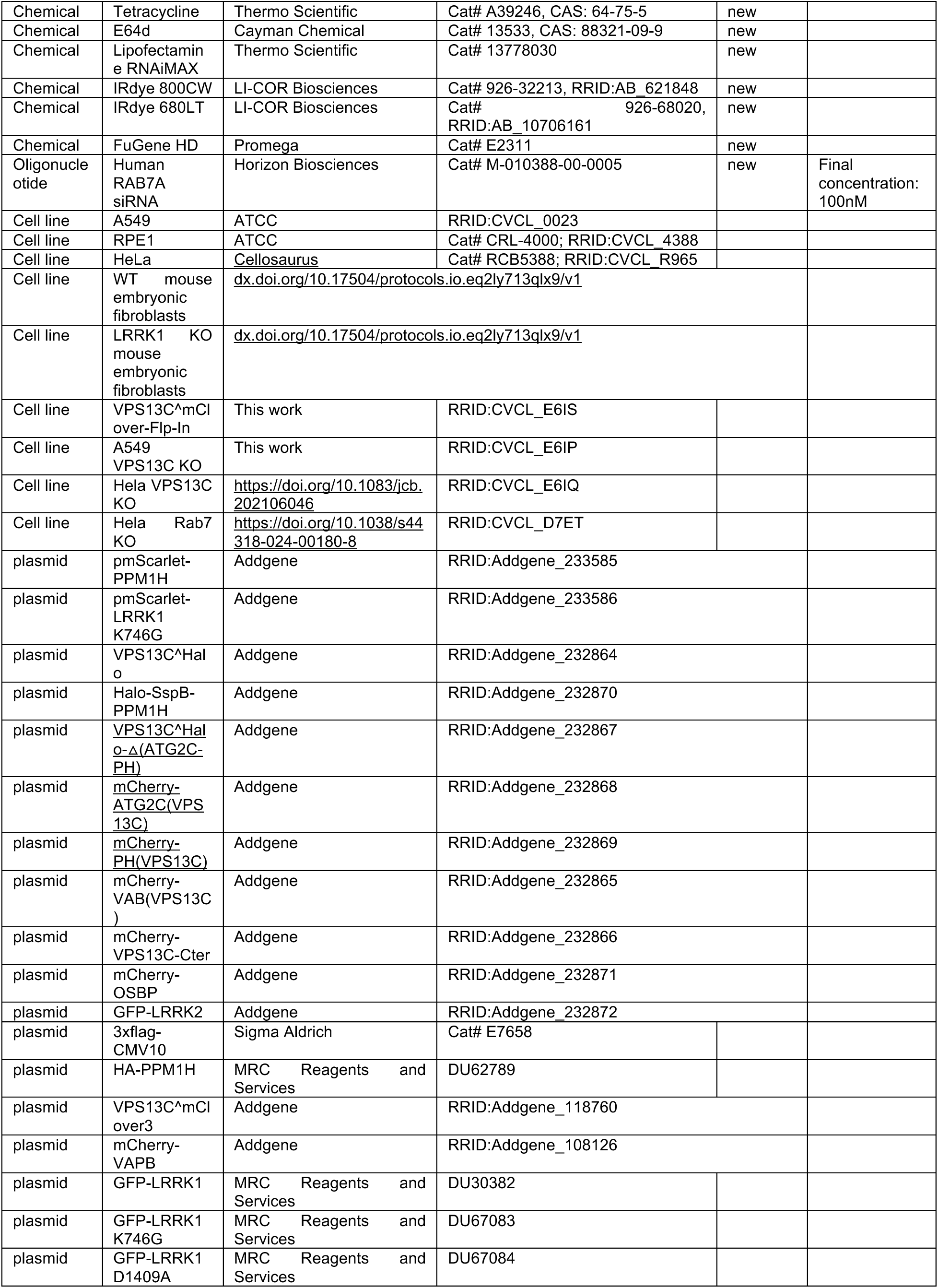
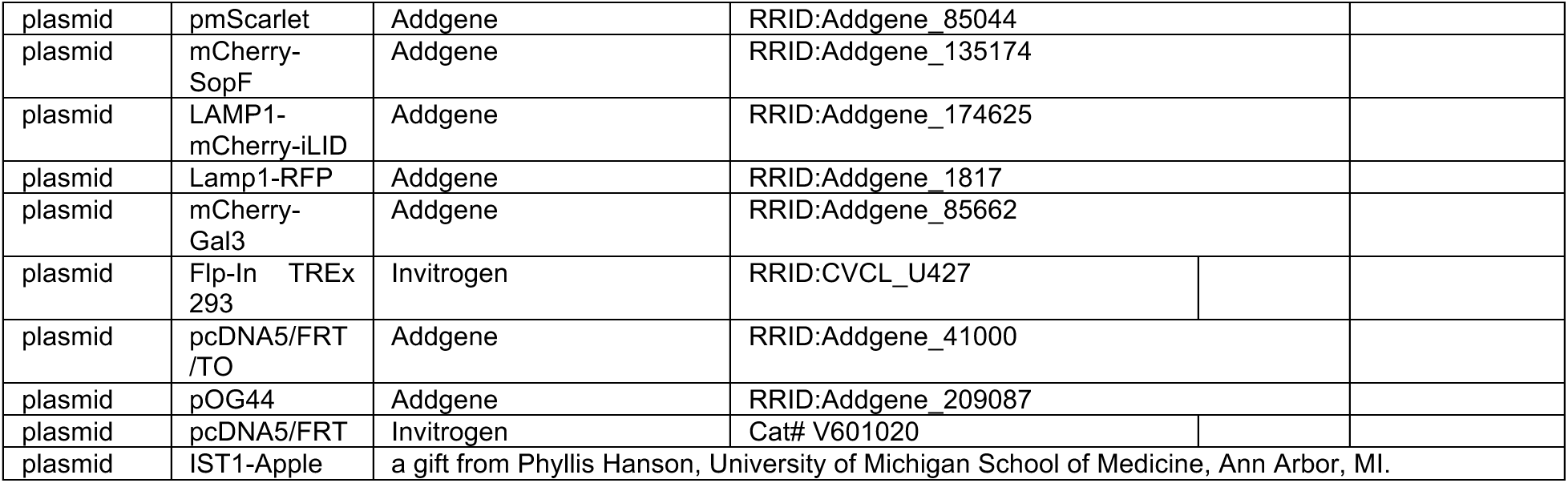

## Notes

### Competing Interest Statement

The authors have declared no competing interest.

